# Oncogenic RAS activity is linked to immune priming and adenosine-driven immune evasion in lung adenocarcinoma

**DOI:** 10.1101/2025.11.25.690426

**Authors:** Sophie de Carné Trécesson, Philip East, Claire E. Pillsbury, Mariana Silva dos Santos, Hongui Cha, Emma Colliver, Tegan Gilmore, Sareena Rana, Chris Moore, Scott Lighterness, Deborah Caswell, Jesse Boumelha, Mona Tomaschko, Romain Baer, Jim Eyles, Beatriz Teixeira, Muhammad Saeed, Kevin Litchfield, Miriam Molina-Arcas, Se-Hoon Lee, James I. MacRae, Philip Hobson, Charles Swanton, Julian Downward

## Abstract

Lung adenocarcinoma (LUAD) is a leading cause of cancer death worldwide, with RAS signalling as a key oncogenic driver. Although *KRAS* mutations have been linked to immune evasion in preclinical models, the relationship between RAS activity and tumour immunity or response to immunotherapy in patients remains unclear. Here, we applied our previously validated RAS84 transcriptional signature to stratify LUAD patient cohorts and dissect the immune landscape associated with RAS signalling. We report that tumours with elevated RAS activity exhibited features of immune priming, including increased immune infiltration, interferon response, and immune checkpoint gene expression, and showed improved progression-free survival in an independent cohort of patients treated with anti-PD-1. Yet, in both LUAD tumours and cell lines, RAS activity also correlated with elevated immunosuppressive interstitial adenosine mediated by transcriptional regulation of several components of the adenosinergic pathway. In orthotopic pre-clinical models of high-RAS activity lung tumours, blocking adenosine signalling delayed tumour growth and improved response to anti-PD-1 and KRAS inhibition, with a significant effect on innate immunity. This study reveals a dual role for RAS signalling in tumour progression, fostering a pro-immunogenic environment whilst simultaneously dampening anti-tumoural immunity via mechanisms including extracellular adenosine accumulation. Stratifying patients based on RAS transcriptional activity, rather than genetic alterations alone, could inform immunotherapy strategies and improve clinical outcomes.

## Introduction

Non-small cell lung cancer (NSCLC) remains the leading cause of cancer-related deaths worldwide^1^. Lung adenocarcinoma (LUAD), its most prevalent subtype, presents a major clinical challenge due to its high mortality and limited treatment efficacy. Immune checkpoint blockade (ICB) targeting the PD-1/PD-L1 axis is now the standard of care for LUAD patients. Anti-PD-1 and anti-PD-L1 therapies offer long-term benefits but only to a subset of patients^2,3^. In contrast, targeted therapy, such as recent KRASG12C inhibitors, has shown excellent response rates, but rapid resistance occurs in virtually all patients receiving those drugs^4^.

A growing number of therapeutic agents designed to reactivate the immune system are entering clinical trials. Notably, agents targeting the adenosine pathway—which leads to immune suppression when activated^5^ —, such as the Adenosine 2A Receptor (A2AR) antagonist ciforadenant (CPI-444) and the anti-CD73 antibody oleclumab, have shown early signs of efficacy in combination with ICB in NSCLC (*e*.*g*., NCT03822351, COAST study; NCT03337698, Morpheus NSCLC)^6,7^. However, clinical responses have been inconsistent, highlighting the need for better patient stratification.

A third of LUAD cases harbour activating mutations in KRAS, making it the most frequently mutated oncogene in this subtype^8^. Beyond its role in promoting cell proliferation and survival, we demonstrated that oncogenic RAS signalling can directly drive immune evasion by increasing PD-L1 expression via the stabilisation of its *CD274* mRNA^9^. Moreover, we and others have shown that inhibiting RAS signalling restores IFN response^10,11^, and reduces expression of the immunosuppressive enzyme COX2 in the cancer cells^12^. Whilst these mechanisms have been characterised in lung and other cancer models, the extent to which RAS signalling influences tumour immunogenicity and response to ICB in LUAD patients remains unclear. Indeed, although some studies have shown a possible association between the presence of KRAS mutations and better outcome in response to anti-PD(L)1, the predictive value of KRAS mutations is ambiguous, with conflicting findings reported across cohorts^13,14^.

We previously demonstrated that RAS pathway activation is not restricted to KRAS-mutant tumours. Using the RAS84 transcriptional signature, we showed that over 80% of LUAD tumours exhibit transcriptional evidence of RAS pathway activity, including KRAS-wild-type tumours^15^. This raises the question of whether RAS pathway activation levels— rather than mutational status alone—better reflect the immune landscape and therapeutic vulnerabilities of LUAD tumours, as we previously showed for chemotherapy response.

Here, we used our previously validated RAS84 classifier to investigate how RAS transcriptional activity shapes the immune microenvironment and modulates response to immunotherapy in LUAD. We reveal that increased levels of RAS activity are associated with T cell infiltration, interferon signalling, immune checkpoint expression, and improved response to anti-PD-1 therapy. However, high-RAS tumours also exhibit elevated extracellular adenosine, a potent immunosuppressive metabolite that inhibits T and NK cell function via A2A receptor signalling^5,16^. We show that adenosine metabolism is directly regulated by RAS signalling to impair the immune response.

This study establishes RAS transcriptional activity as a key determinant of the immune landscape and the sensitivity to immunotherapy in LUAD. Our findings reveal a dual role for RAS signalling, promoting tumour immunogenicity whilst driving immune evasion through activation of the adenosine pathway. These insights provide a rationale for combining ICB with adenosine pathway inhibition in high-RAS tumours, and position RAS84 as a potential biomarker to personalise immunotherapy strategies in LUAD.

## Results

### Oncogenic RAS signalling correlates with inflammation and immune checkpoint gene expression

To understand how oncogenic RAS influences the immune response in human lung cancer, we used the RAS84 signature and classification of TCGA lung adenocarcinoma (LUAD) we previously established^15^. This method uses RAS transcriptional activity as a surrogate for oncogenic RAS activity. Hence, it considers any alterations other than the KRAS mutation that can affect RAS pathway activity (*e*.*g*., *NF1* or *EGFR* mutation, *ALK* fusion, *KRAS* amplification). We classified 517 LUAD samples in 5 RAS Activity Groups, hereafter named RAGs, with increasing RAS-index (RI), from RAG-0 (with an RI close to zero) to RAG-4 (highest RI) (**Figure 1a**). RAS84 was designed and tested to quantify RAS activity in the cancer cells, with minimal signal in the immune compartment^15^. To untangle the effect of RAS signalling on the immune landscape from its autonomous effect, we classified lung cell lines from the CCLE dataset^17^ (**Supp. Data Table 1**).

**Figure 1.**
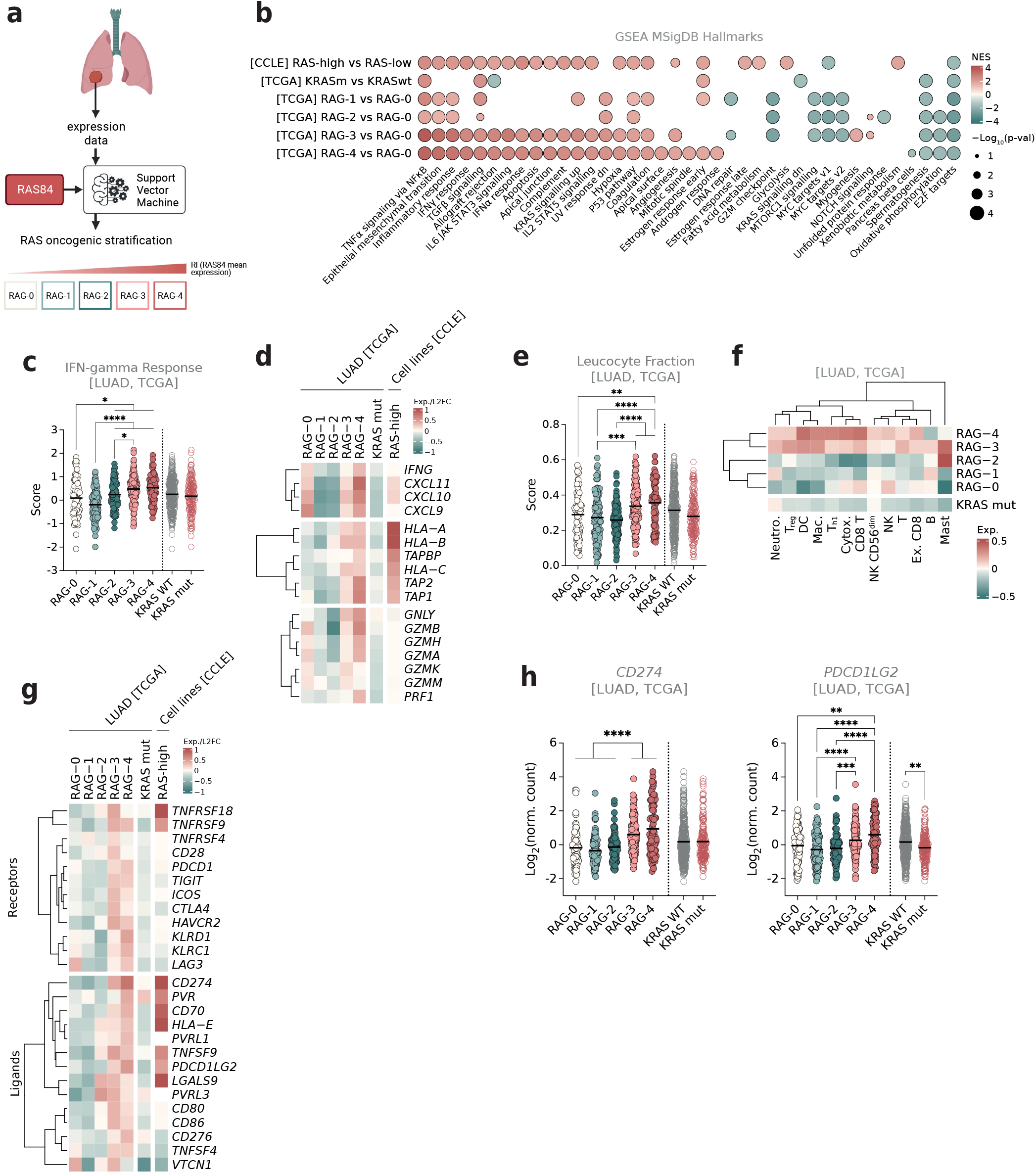
RAS oncogenic activity is associated with inflammation and immune-checkpoint gene expression. (a) Schematic showing the method used to classify samples according to their level of RAS transcriptional activity from East et al. (2022)^15^. (b) Normalised enrichment scores (NES) and p-values from GSEA of enriched HALLMARK genesets in each RAG compared with RAG-0 and KRAS mutants compared with WT in LUAD tumours (TCGA), and RAS-high versus RAS-low lung adenocarcinoma cell lines (CCLE). Only the significant NES are shown (FWER p-val < 0.05). (c) IFN-gamma response score from Thorsson et al. (2018)^18^ mean and individual expression per RAG or KRAS mut and WT groups in LUAD tumours (TCGA). Brown-Forsythe and Welch ANOVA tests corrected for multiple comparisons using Games-Howell’s tests (*p≤0.05, **p≤0.01, ****p≤0.0001). (d) Heatmap showing the mean expression of immune-related genes scaled to median gene values across RAGs or log2FC between KRAS mutant and WT groups in LUAD (TCGA) or between RAS-high and RAS-low lung adenocarcinoma cell lines (CCLE). (e) Leucocyte fraction score calculated from methylation data (see Thorsson et al. 2018)^18^, mean and individual expression per RAG or KRAS mut and WT groups in LUAD tumours (TCGA). Brown-Forsythe and Welch ANOVA tests corrected for multiple comparisons using Games-Howell’s tests (*p≤0.05, **p≤0.01, ***p≤0.001, ****p≤0.0001). (f) Heatmap showing immune cell scores (adapted from Danaher et al. 2017)^19^ scaled to median gene signature values across RAGs or log2FC between KRAS mutant and WT groups in LUAD (TCGA). (g) Heatmap showing the mean expression of immune checkpoint receptors (top) or ligands (bottom) genes scaled to median gene values across RAGs or log2FC between KRAS mutant and WT groups in LUAD (TCGA) or between RAS-high and RAS-low lung adenocarcinoma cell lines (CCLE). (h) *CD274* and *PDCD1LG2* mean and individual expression per RAG or KRAS mut and WT groups in LUAD tumours (TCGA). Brown-Forsythe and Welch ANOVA tests corrected for multiple comparisons using Games-Howell’s tests (**p≤0.01, ***p≤0.001, ****p≤0.0001).

Gene-Set Enrichment Analysis (GSEA) revealed a significant enrichment of several genesets associated with an immune response in RAG-3 and -4 (*e*.*g*., TNF*α* signalling via NF*κ*B, inflammatory response, IFN*γ* response, allograft rejection, IFN*α* response, IL2-STAT5 signalling, and Complement, also enriched in cell lines (**Figure 1b, Supp. Data Table 2**). As expected, processes known to be regulated by RAS were also enriched in these groups and not in KRAS mutant (KRASm) compared with KRAS wild type (KRASwt) (*e*.*g*., KRAS signalling, EMT, angiogenesis), confirming the superior power of RAS84 to evaluate RAS signalling compared with the mutational status of KRAS only.

The IFN*γ* response signature score was particularly low in RAG-1 (**Figure 1c**). Th1 chemoattractant cytokines (*CXCL10, CXCL11*, and *CXCL9*), antigen-presenting machinery genes (*TAP1, TAP2, HLA-A, HLA-B, HLA-C*, and *B2M*), and cytotoxic enzymes (PRF1, GZMA, and GZMB) were all highly expressed in RAG-3 and -4 compared with RAG-1 and -2 tumours (**Figure 1d**), suggesting the recruitment of cytotoxic cells and the possibility of presenting neoantigens in high-RAS-driven tumours.

Using the leucocyte score previously calculated by Thorson and colleagues from methylation data in this cohort^18^, we found an increase in leucocyte fraction in RAG-3 and -4 compared with other groups, particularly when compared with RAG-1 and -2 (**Figure 1e**). We confirmed this finding using the RNA-Seq data by looking at the expression of the Danaher and Davoli immune signatures^19,20^, previously validated by Rosenthal and colleagues^21^ (**Figure 1f, Supp. Data Table 3** and **Supp. Figure 1a**). RAG-3 and -4 were highly infiltrated with CD4 and CD8 T cells (including Th1 and exhausted CD8), macrophages, dendritic cells, and neutrophils. In contrast, RAG-2 tumours exhibited low levels of CD8 T, cytotoxic cells, including NK cells, and high levels of mast cells, which were particularly specific to this group. RAG-3 had elevated Treg levels. These results suggest that increased levels of oncogenic RAS activity are associated with high immune infiltration. Interestingly, neoantigen counts were lower in RAG-1 to -4 compared with RAG-0, suggesting that our observations were not simply due to increased antigen presentation in RAS-high tumours (**Supp. Figure 1b**). Notably, RAG-0 patients exhibited a significant increase in the proportion of tumours with loss of heterozygosity (LOH) of at least one HLA gene in the TRACERx LUAD cohort (39%, whilst RAG-1 to -4 showed 10%, 22%, 23%, and 32% of HLA LOH, respectively), indicating a potential reduction in neoantigen presentation in this group (**Supp. Figure 1c**). We previously showed that RAG-3 and RAG-4 patients had a worse outcome than RAG-0 resectable LUAD^15^. Given the high infiltration of Th1 and exhausted CD8 T cells in these groups, we examined the expression of immune checkpoint receptors and ligands to understand how these tumours evade immune responses. As expected, the increased lymphocyte presence coincided with higher expression levels of most immune checkpoint receptors and ligands in RAG-3 and RAG-4 compared with RAG-0 to -2 (**Figure 1g**), notably CD274 (PD-L1) and PDCD1LG2 (PD-L2) (**Figure 1h, Supp. Data Table 4 and 5**).

This analysis illustrates that RAG-3 and RAG-4 tumours are immune-infiltrated and have a high IFN*γ* response. It suggests that high RAS84 expression is associated with an anti-tumoural immune response likely dampened by the expression of immune checkpoint ligands such as *CD274* and *PDCD1LG2*. Altogether, it suggests that RAG-3 and RAG-4 tumours are primed to respond to immune checkpoint blockade (ICB).

### Oncogenic RAS transcriptional activity is associated with better response to ICB

To test whether RAG-3 and RAG-4 tumours responded better to ICB, we investigated a cohort of 314 LUAD patients from the Samsung Medical Centre in South Korea^22^. All patients received anti-PD-1 treatment and showed varying levels of RECIST response; however, no patients achieved complete response. We stratified the patients by applying our classifier to the RNA-Seq data from tumour biopsies before treatment^15^ (**Supp. Figure 2a**). RAG-3 and RAG-4 had the most prolonged median PFS, respectively, 5.27 and 3.80 months (**Figure 2a**). Forty-one per cent of patients in these groups were partial responders (PR) (**Figure 2b**). In contrast, RAG-2 patients had the worst outcome (median PFS, 1.83 months), and only one patient (2%) achieved PR in this group. In comparison, KRAS-mutant patients did not have a significantly improved PFS compared with KRAS WT patients, demonstrating that RAS transcriptional activity outperforms KRAS mutational status in predicting response to ICB (**Supp. Figure 2b**). Multivariate Cox proportional Hazards Analysis showed that RAG-3, and RAG-4 patients had significantly better PFS when compared with RAG-2 patients (**Figure 2c**). As observed in the TCGA LUAD cohort, *CD274* mean expression was markedly higher in RAG-3 and RAG-4 compared with the other groups in the Samsung cohort (**Figure 2d**).

**Figure 2.**
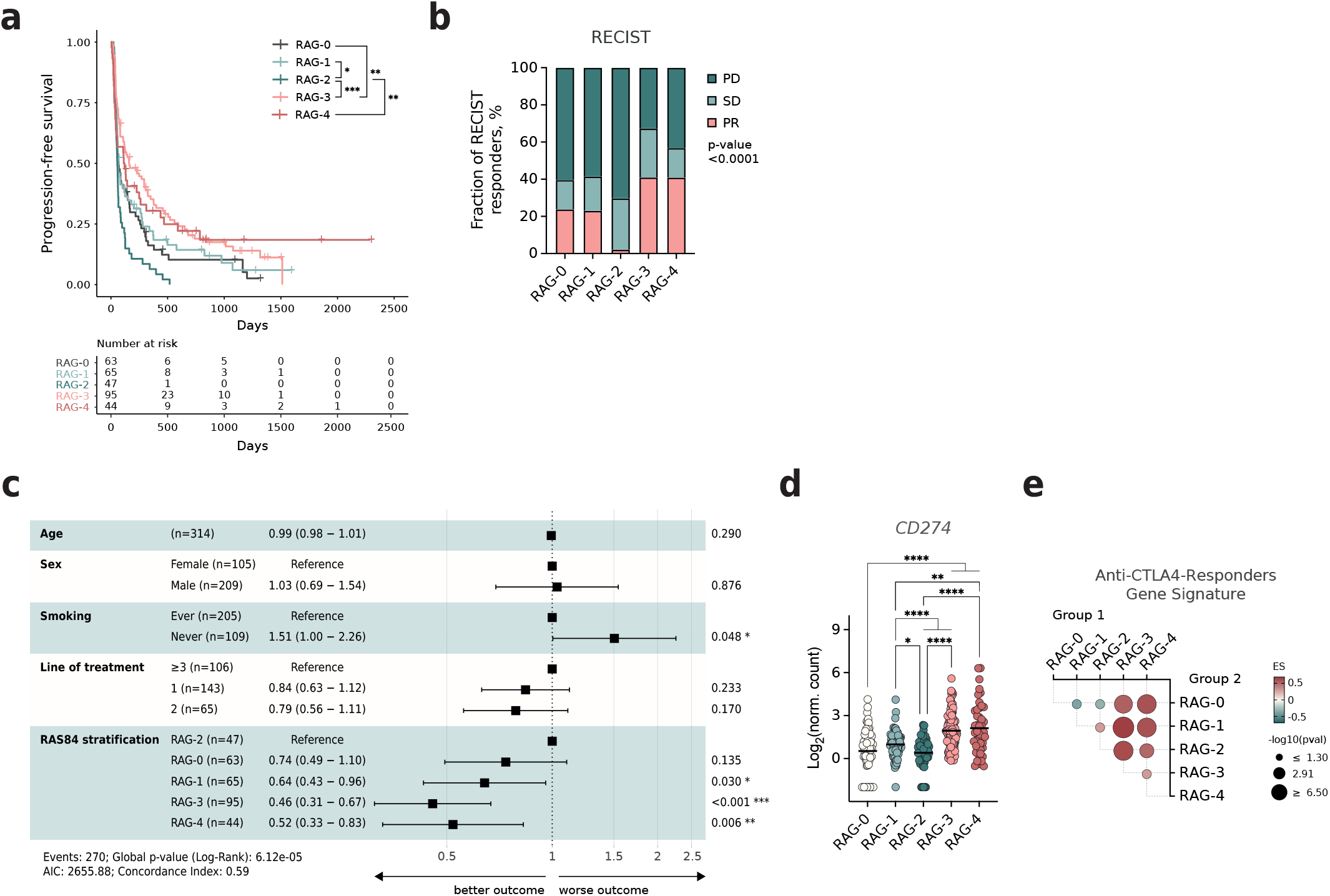
RAS oncogenic activity predicts response to anti-PD-1 in LUAD. (a) Kaplan-Meier estimate of the progression-free survival in LUAD patients from the Samsung Medical Center in South Korea stratified with RAS84 (n = 314 patients). (b) Fraction of RECIST patients per RAG (Samsung cohort). Chi-square test (p = 1.5×10^−5^). (c) Forest plot of hazard ratios and corresponding 95% confidence intervals estimated from multivariate Cox models for overall progression-free survival in relation to RAG stratification. All LUAD patients received anti-PD-1 treatment. Samsung cohort (n = 314 LUAD patients). (d) *CD274* mean and individual expression in pre-treatment tumour biopsy per RAG in LUAD tumours (Samsung cohort). Brown-Forsythe and Welch ANOVA tests corrected for multiple comparisons using Games-Howell’s tests (**p≤0.01, ***p≤0.001, ****p≤0.0001). (e) Enrichment score (ES) and p-value from GSEA of anti-CTLA-4-responders gene signature (Ock et al. 2017)^23^ in LUAD (TCGA) RAGs. Comparisons are shown as group 1 versus group 2. Only the significant ESs are shown (p≤0.05).

To extend the observation to another immune checkpoint inhibitor, we performed a GSEA of a gene signature associated with anti-CTLA-4 responders developed by Ock and colleagues^23^. Anti-CLTA-4-responder genes were systematically enriched in RAG-3 and RAG-4 in all comparisons (**Figure 2e** and **Supp. Figure 2c**).

Together, these findings demonstrate that RAS84 stratification predicts benefit from immune checkpoint blockade more accurately than KRAS mutation status, positioning RAG-3 and RAG-4 tumours as the most immunologically responsive LUAD subtypes.

### Oncogenic RAS activity correlates with NT5E and increased levels of extracellular adenosine

Despite showing the best responses to ICB, only 40% of RAG-3 and RAG-4 patients achieved partial responses, and no complete responses were observed in this cohort. This suggests other immune evasion mechanisms likely blunt immunotherapy efficacy in primed tumours. To identify such mechanisms potentially driven by RAS signalling, we reasoned that relevant genes would correlate with RAS transcriptional activity across tumours. We thus calculated the Pearson correlation coefficient between each gene and RAS84 expression across LUAD tumours and NSCLC cell lines (**Figure 3a, Supp. Data Table 6**). As expected, RAS target genes—including RAS84 genes—had high Pearson correlation coefficients in both datasets. Among the most correlated genes, we also identified *NT5E*, which codes for CD73, an ectonucleotidase that converts extracellular AMP into adenosine, a potent inhibitor of the immune response^5^. Several genes coding for proteins involved in the metabolism of interstitial adenosine (iADO) metabolism were differentially expressed across RAGs of LUAD tumours and cell lines (**Figure 3b**). CD73 protein expression was highly correlated with *NT5E* mRNA expression (Pearson correlation coefficient = 0.86, p-value <0.0001) across cell lines, suggesting that CD73 expression in cancer cells is primarily regulated transcriptionally (**Figure 3c**). *NT5E* and *ADORA2B*, a receptor for iADO expressed by epithelial cells, were overexpressed in RAG-2, -3, and -4 compared with RAG-0 and -1 (**Figure 3d**). In contrast, *NT5E* expression was only moderately increased in KRAS mutants versus KRAS WT tumours (L2FC 0.42, padj≤0.05), and no other genes in the adenosine pathway showed significant differential expression between these groups (**Supp. Figure 3b**).

**Figure 3.**
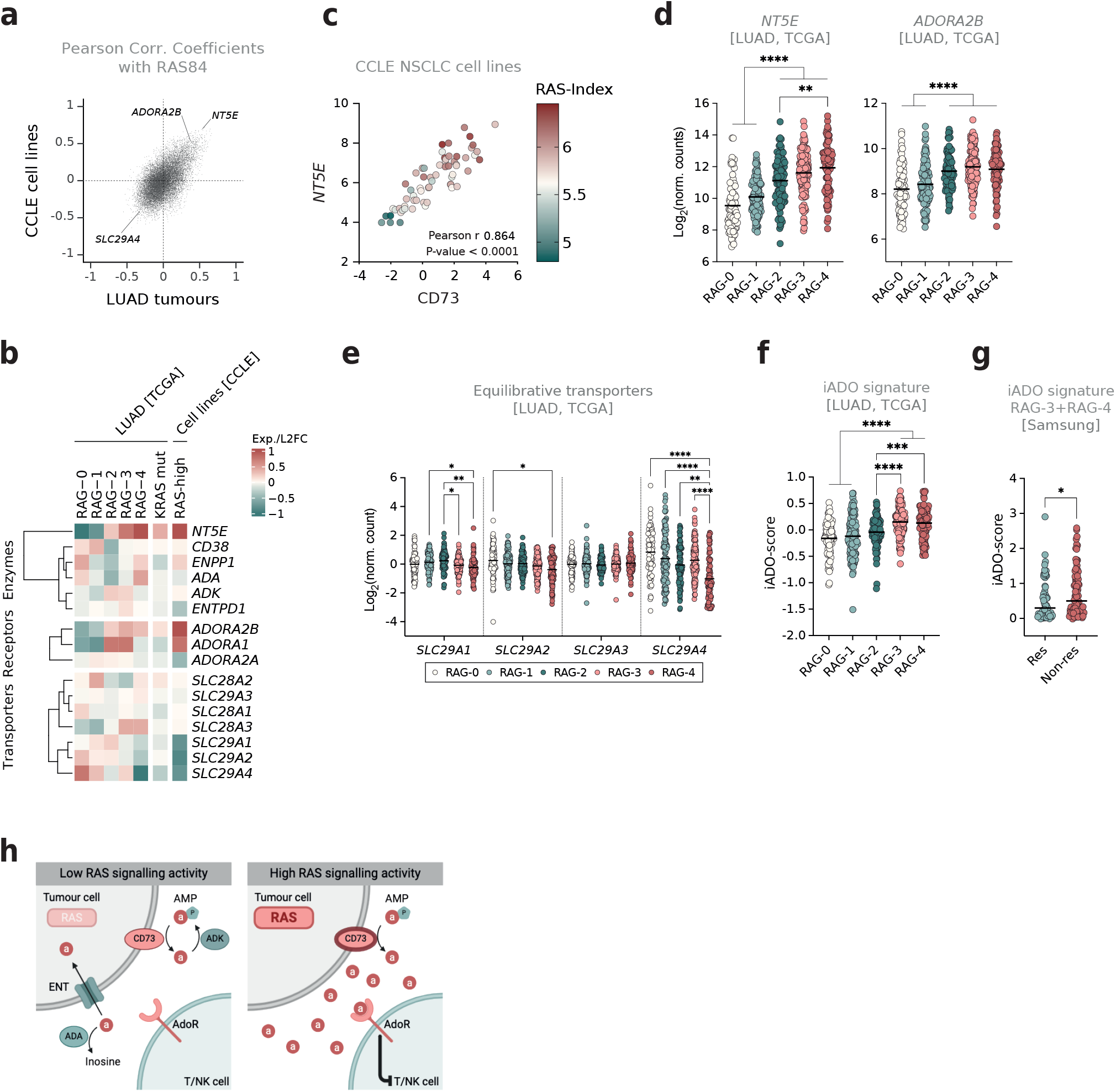
Increased RAS oncogenic activity is associated with high levels of extracellular aden-osine. (a) Scatterplot showing Pearson correlation coefficients between the RAS84-index and gene expression levels for all genes across CCLE NSCLC cell lines (n = 63) (y-axis) and TCGA LUAD patient tumours (n = 502) (x-axis). *NT5E, ADORA2B* and *SLC29A4* are indicated. (b) Heatmap showing the mean expression of adenosine-metabolism-related genes scaled to median gene values across RAGs or log2FC between KRAS mutant and WT groups in LUAD (TCGA) or between RAS-high and RAS-low lung adenocarcinoma cell lines (CCLE). (c) Scatterplot showing relative expressions of *NT5E* mRNA (y-axis), its protein CD73 (x-axis), and the RAS-index (colour scale) of all CCLE NSCLC cell lines with both data available (n = 63). (d) *NT5E, ADORA2B*, and *SLC29A4* mean and individual expression per RAG in LUAD tumours (TCGA). Brown-Forsythe and Welch ANOVA tests corrected for multiple comparisons using Games-Howell’s tests (*p≤0.05, **p≤0.01, ***p≤0.001, ****p≤0.0001). (e) Equilibrative transporter genes presented as mean per group (RAG) and individual tumour values in LUAD (TCGA). Brown-Forsythe and Welch ANOVA tests corrected for multiple comparisons using Games-Howell’s tests (*p≤0.05, **p≤0.01, ****p≤0.0001). (f) Interstitial adenosine signature score (iADO-score) (Sidders et al. 2019)^25^ mean and individual expression per RAG in LUAD tumours (TCGA). Brown-Forsythe and Welch ANOVA tests corrected for multiple comparisons using Games-Howell’s tests (***p≤0.001, ****p≤0.0001). (g) Interstitial adenosine signature score (iADO-score) (Sidders et al. 2019)^25^ median and individual expression per RAG in LUAD tumours (Samsung). Mann-Whitney U test (*one-tailed p=0.0283). (h) Schematic showing the regulation of interstitial adenosine in the tumour environment and its effect on T and NK cells. “a” in red circles = adenosine metabolite, “P” in the blue heptamer = phosphate group, ENT = equilibrative nucleotide transporters, AdoR = adenosine receptor.

Equilibrative nucleoside transporters (ENTs) allow the passive diffusion of nucleosides along their concentration gradient, whereas concentrative nucleoside transporters (CNTs) are Na^+^-dependent active transporters that transport nucleosides into cells^24^. ENT genes were expressed at higher levels in LUAD tumours (**Supp. Figure 3c**), and *SLC29A1, SLC29A2*, and *SLC29A4* were all downregulated in RAG-4, with *SLC29A4* having the largest effect size compared with RAG-0 (L2FC −1.99) (**Figure 3e**). The increase in *NT5E* and decrease in ENT gene expression suggest the accumulation of iADO in the extracellular space in tumours, which was corroborated by the significant increase in iADO-scores^25^ in RAG-3 and RAG-4 tumours compared with RAG-0 to -3 (**Figure 3e**).

To assess whether iADO could prevent a potent immune response in immune-primed tumours, we compared iADO scores in RAG-3 or RAG-4 responders and non-responders to anti-PD-1 immunotherapy in the Samsung cohort. Compared with non-responders, responder tumours were notably enriched in *CD8A, CD274, EOMES, TBX21, IFNG*, and *PDCD1*, suggesting the presence of exhausted T cells (**Supp. Figure 3d**), and iADO scores were significantly higher in the non-responders (**Figure 3f**). The mean expression of *NT5E* increased incrementally in Samsung RAGs, as observed in the TCGA LUAD cohort (**Supp. Figure 3e**). Of note, iADO scores were not different in KRAS mutant and WT tumours (**Supp. Figure 3f**).

Altogether, these data suggest that RAS activity is associated with changes in adenosine metabolism, which could impair T cell response and promote resistance to anti-PD-1 therapy (**Figure 3h**).

### Oncogenic RAS signalling regulates extracellular adenosine levels in vitro

We then investigated whether modulating RAS signalling could influence adenosine metabolism gene expression. We used three lung cancer cell lines harbouring the KRAS^G12C^ mutation: H23, H1792, and H358. Treatment with a KRASG12C inhibitor (G12Ci) for 24 hours decreased the expression of *NT5E* and *ADORA2B* whilst inducing the expression of *SLC29A4* and *ADA* (adenosine deaminase) across all cell lines. Additionally, *ADK* (adenosine kinase) expression was induced in the H23 and H358 cell lines (**Figure 4a**). ADA catalyses the conversion of adenosine to inosine, while ADK catalyses the phosphorylation of adenosine to AMP, thereby reducing adenosine levels in the tumour. *SLC29A1* expression was altered only in H358, where G12Ci decreased its expression. The baseline expression profiles of adenosine-related genes across the three cell lines suggested potential co-regulation of these genes (**Supp. Figure 4**). Indeed, we observed strong positive and negative correlations between *NT5E, ADORA2B, ADA*, and *SLC29A4* across samples, along with a negative correlation between *ADK* and *SLC29A1* expression, supporting the hypothesis of their co-regulation (**Supp. Figure 4b**). The regulation of *NT5E* expression by G12Ci also occurred in the colon cancer cell line SW873, suggesting that RAS signalling promoting interstitial adenosine accumulation may be generalised to other cancers (**Supp. Figure 4c**). We next assessed CD73 protein expression following RAS signalling inhibition. A significant reduction in CD73 protein levels required 72 hours of G12Ci treatment, indicating a long protein half-life (**Figure 4b**). We confirmed this reduction in CD73 expression at the plasma membrane after 72 hours of treatment (**Supp. Figure 4d**). These findings suggest that inhibiting RAS signalling could prevent the accumulation of extracellular adenosine.

**Figure 4.**
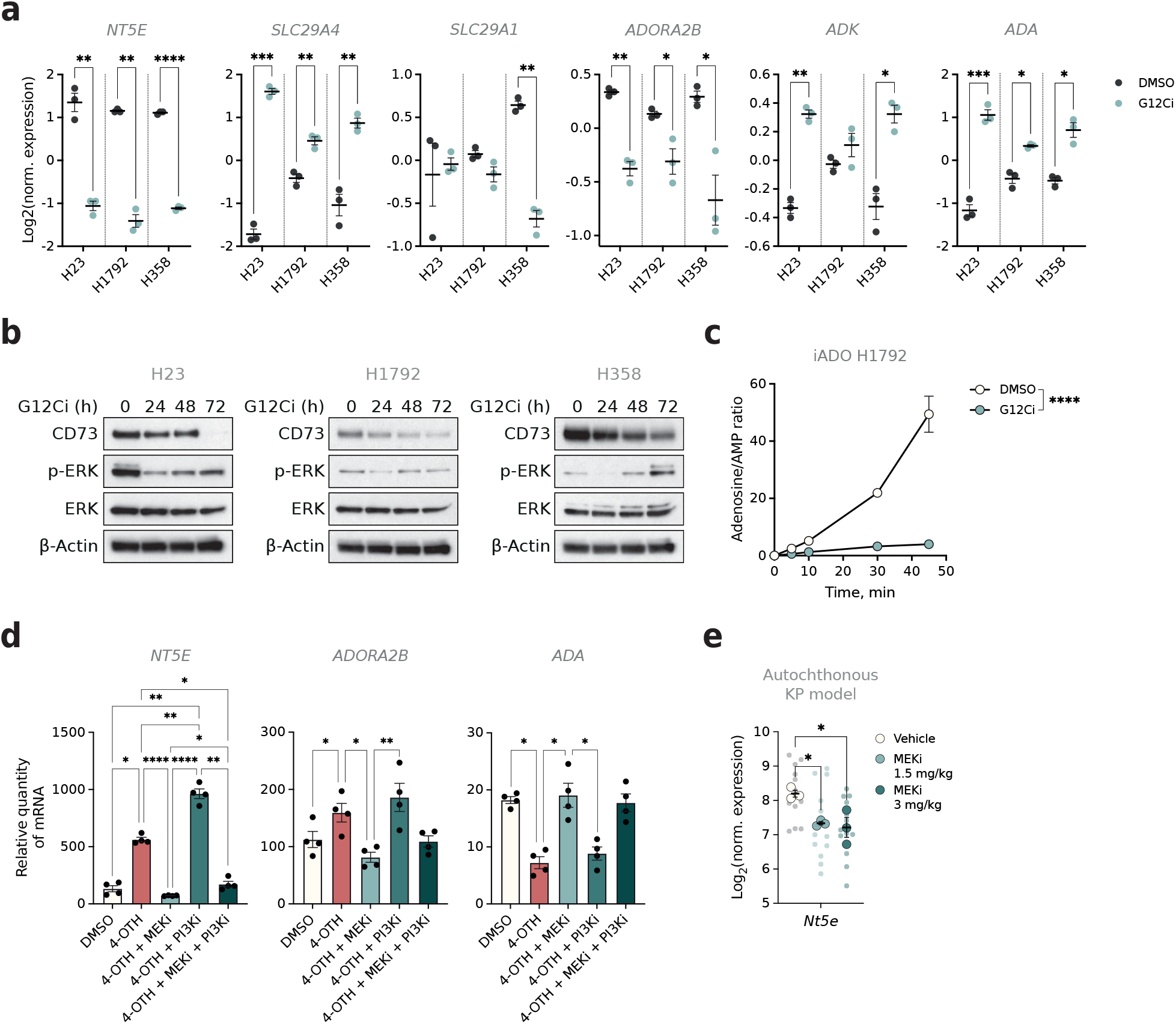
RAS activity modulates adenosine-related genes and extracellular AMP-to-adenosine metabolisation. (a) *NT5E, SLC29A4, SLC29A1, ADORA2B, ADK* and *ADA* expression from qPCR, mean ±SEM per treatment group, dots represent independent biological replicates (n = 3). The indicated cell lines received DMSO or 10 *μ*M ARS853 for 24h. One-sided p-value with unequal variances *t*-test performed between treatment groups (*p≤0.05, **p≤0.01, ***p≤0.001, ****p≤0.0001). (b) Western blot showing CD73, p-ERK1/2 (Thr202/Tyr204), ERK1/2 and B-actin in human lung cancer cell lines treated with 10 *μ*M ARS1620 for the indicated time. (c) Adenosine-to-AMP ratio calculated from mass spectrometry analysis of the supernatant of H1792 cells pre-treated with DMSO or 200 nM MRTX1257 for 72h. 100 nM AMP was added to the supernatants at time 0. Mean ±SD of 4 technical replicates per time and treatment group. RM two-way ANOVA with the Geisser-Greenhouse correction, Šidák’s multiple comparisons test (****p≤0.0001). (d) *NT5E, ADORA2B*, and *ADA* expression in the AT2-KRAS^G12V^ cell line from qPCR, mean ±SEM per treatment group, dots represent independent biological replicates (n = 4). KRAS^G12V^ was induced with 100 nM 4-OTH for 24h in the presence or not of 10 nM trametinib (MEKi) and/or 500 nM GDC0941 (PI3Ki). One-way ANOVA, with the Geisser-Greenhouse correction. Tukey’s multiple comparisons test (*p≤0.05, **p≤0.01, ***p≤0.001, ****p≤0.0001). (e) *Nt5e, Cd3*, and *Ncr1* expression from qPCR, mean ±SEM per treatment group. Large dots represent the mean per mouse (n = 3 per group), and small dots represent individual tumour expression. KP-tumour-bearing mice received 4 doses of trametinib over 3 days. Brown-Forsythe and Welch ANOVA tests performed on mean per mouse, corrected for multiple comparisons using Games-Howell’s tests (ns p>0.05, *p≤0.05).

To test this hypothesis, we treated H1792 cells with G12Ci for 72 hours to downregulate CD73 expression. We chose H1792 cells because H23 and H358 cell lines are sensitive to G12Ci and induce cell death upon treatment^26^. At 72h, cell proliferation and viability were moderately affected by RAS signalling inhibition in H1792 cells (**Supp. Figure 4e**). We then added AMP to the supernatant and measured adenosine and AMP levels over time. Whilst AMP was rapidly metabolised into adenosine in the supernatant of untreated cells, this process was abrogated in G12Ci-treated cells (**Figure 4c**), supporting the hypothesis that RAS signalling promotes the increase in interstitial adenosine by regulating the expression of CD73 and several other adenosine-related genes.

We tested the effect of activating RAS signalling using a previously described model of immortalised human lung pneumocytes expressing a tamoxifen-inducible KRAS^G12V^ (KRAS^G12V-ER^)^27^. KRAS^G12V^ activation induced *NT5E* and *ADORA2B* and repressed *ADA* expression. Notably, these effects were fully reversed by MEK inhibition, but not by PI3K inhibition, suggesting that the RAS–MEK–ERK pathway is primarily responsible for regulating these adenosine metabolism genes (**Figure 4d** and **Supp. Figure 4f**).

We validated the relevance of this regulatory axis by treating *Kras*^G12D-LSL/WT^; *Trp53*^fl/fl^ (KP) tumour-bearing mice with MEK inhibitor. *Nt5e* expression was significantly decreased following MEKi treatment, confirming the role of MAPK in the regulation of this pathway *in vivo* (**Figure 4e**).

Altogether, these data show that oncogenic RAS signalling promotes the accumulation of extracellular adenosine by regulating several genes that code for proteins involved in adenosine metabolism.

### Oncogenic RAS signalling is associated with immune priming in mouse lung tumours

The KP tumours mentioned above are considered ‘immune cold’ due to the absence of neoantigens presented by the cancer cells, consequent to low genomic mutation load. To better model LUAD, and particularly immune-primed tumours with high RAS signalling, we used KPAR and 3LL murine orthotopic transplant models. The 3LL cell line originates from a Lewis lung carcinoma, carries a *Kras*^G12C^ mutation, and is highly immune-evasive despite having a high mutational load^10^. The KPAR cells were established in our lab from a KP tumour expressing human APOBEC3B in a *Rag1*^KO/KO^ background and exist in two versions: KPAR1.3 with a *Kras*^G12D^ mutation and KPAR^G12C^ with a *Kras*^G12C^ mutation^28^. We previously showed that KPAR tumours respond partially to ICB, whereas 3LL tumours are refractory to ICB^10,28^.

To estimate the levels of RAS signalling in these mouse models, we translated the RAS84 signature to its murine version, mRAS79 (**Supp. Data Table 7**). Using RNA-Seq data, we estimated RAS signalling activity in both cell lines. KPAR cells had a higher mRAS79-index than 3LL cells when cultivated *in vitro* (**Figure 5a**). We then applied a linear regression model to the mRAS79 gene expression data from RNA-Seq analyses of 3LL and KPAR lung tumours to account for species differences and integrate the mouse tumours into our human LUAD RAG classifier. PCA analysis revealed that KPAR tumours clustered closer to RAG-4, whereas 3LL tumours were closer to RAG-1 (**Figure 5b**). This observation was confirmed by clustering the tumour samples using the RAS84 genes that drove the clustering of human RAGs (**Supp. Figure 5a**).

**Figure 5.**
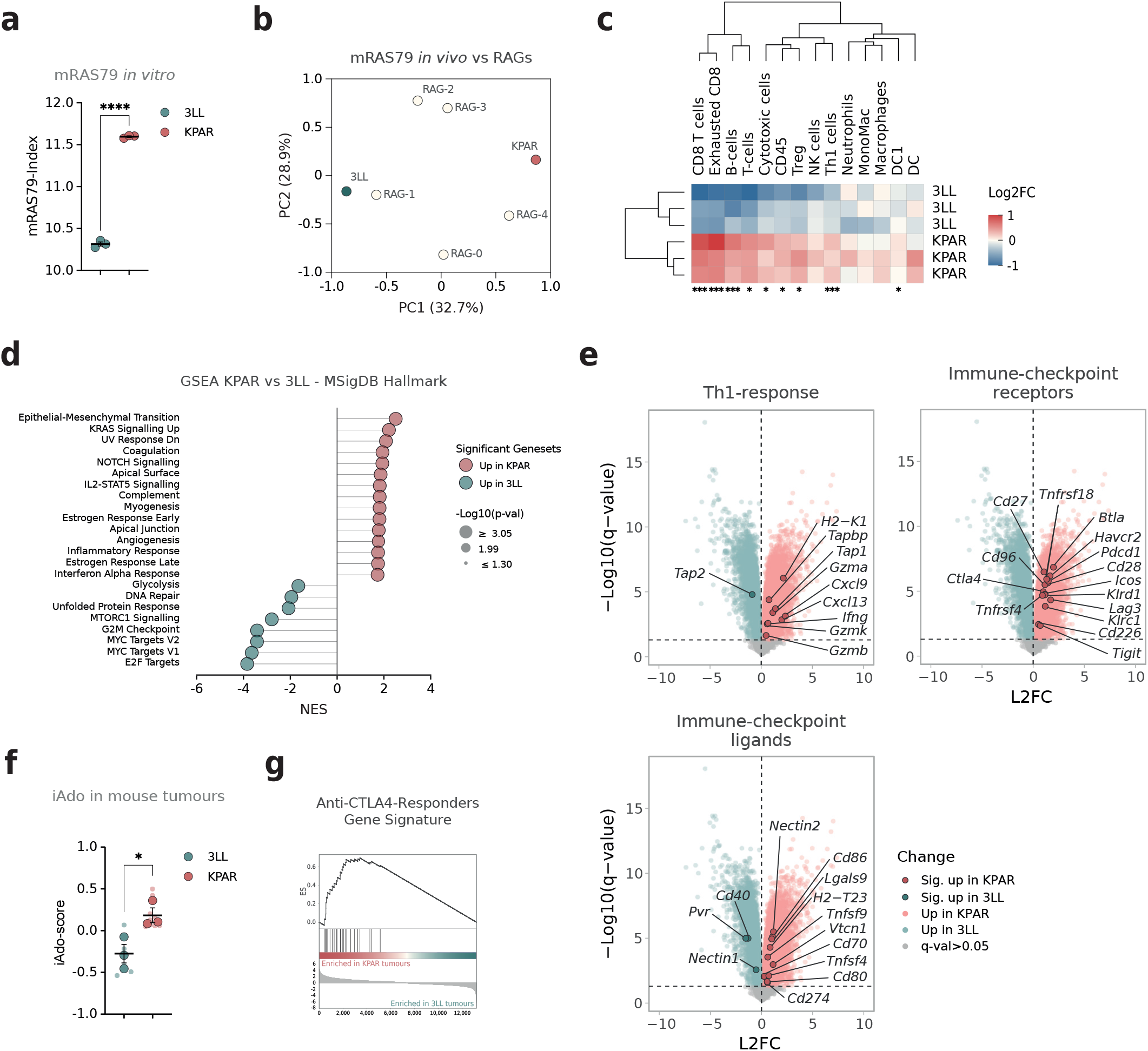
Modelling RAGs in pre-clinical experiments. (a) mRAS79 expression form RNA-Seq data in 3LL and KPAR cell lines cultivated *in vitro*, mean and ±SEM (n = 3 independent experiments). Mann-Whitney U test (****one-tailed p≤0.0001). (b) Principal Component Analysis (PCA) of integrated mouse 3LL and KPAR tumour models with human RAG tumours using mRAS79. Axis lengths are representative of the proportion of variance reported in brackets for PC1 and PC2. (c) Heatmap showing immune cell scores (adapted from Danaher et al. 2017)^19^ scaled to median gene values across samples in 3LL and KPAR tumours. Unpaired *t*-test with Welch’s correction, followed by a two-stage linear step-up procedure (Benjamini, Krieger, and Yekutieli) for false discovery rate (FDR) control. P-values are reported where FDR ≤ 0.05. (d) Interstitial adenosine signature score (iADO-score) (Sidders et al. 2019)^25^ mean per model ±SEM (n = 3 mice per model). Large dots represent the mean score per mouse, and the small dots represent individual tumour scores. Unpaired t-test (*one-tailed p≤0.05). (e) Enrichment plot of anti-CTLA-4-responders gene signature (Ock et al. 2017)^23^ from GSEA between 3LL and KPAR tumours. (f) Normalised enrichment score (NES) and p-value from GSEA of enriched HALLMARK genesets KPAR (n = 9 tumours from 3 mice) versus 3LL (n = 6 tumours from 3 mice) tumours. Only the significant NES are shown (FWER p-val ≤ 0.05). (g) Volcano plots showing Differential Gene Expression between KPAR and 3LL tumours, Log2FC and −log10(p-value). Unchanged genes in grey are genes with −log10(p-value) <1.3, equivalent to a p-value >0.05. The annotated genes correspond to the significantly differentially expressed genes presented in Fig.1.

In line with the high immune infiltrate observed in RAG-4, KPAR tumours were enriched in T cells, including Cd8 and exhausted Cd8, B cells, cytotoxic cells, Th1, Treg, cDC1, and Cd45^+^ cells in general, compared with 3LL tumours (**Figure 5c**). Interestingly, the mRAS79-index (RI) correlated with the presence of exhausted Cd8^+^ cells across tumours within both models, suggesting that increased RAS activity promotes Cd8^+^ T cell exhaustion, even in poorly infiltrated tumours (**Supp. Figure 5b** and **c**). In addition, RI positively correlated with T cell infiltration in 3LL tumours and negatively correlated with macrophage abundance in both models and dendritic cell presence in KPAR tumours.

GSEA revealed that several immune-related gene sets, such as Complement, Inflammatory Response, and Interferon Alpha Response, were enriched in KPAR tumours compared with 3LL tumours. In contrast, 3LL tumours were enriched in Myc Target Genes V1 and V2, consistent with their immune-excluded status. Additionally, KRAS Signalling Up genes were enriched in KPAR tumours compared to 3LL tumours, further validating that KPAR tumours display higher RAS signalling activity than 3LL tumours (**Figure 5d**). Genes associated with a type 1 helper T cell (Th1) response, including *Ifng* itself, were overexpressed in KPAR tumours, as were immune checkpoint receptors and ligands, except for *Cd40, Pvr*, and *Nectin1*, which were overexpressed in 3LL tumours. The latter two regulate T or NK cell inhibition through binding to the Tigit receptor (**Figure 5e**).

Finally, KPAR tumours expressed higher iAdo-scores than 3LL (**Figure 5f**) and were enriched with the anti-CTLA-4 responder gene signature. Altogether, this analysis suggests that KPAR tumours replicate the characteristics observed in RAG-4 tumours in terms of RAS signalling, immune infiltration, and response to ICB, whereas 3LL tumours recapitulate RAG-1 or perhaps RAG-2 tumours, characterised by a low immune infiltrate and resistance to ICB. Despite these immune-primed features, KPAR tumours also display a strong adenosine signature, suggesting that extracellular adenosine may act as a limiting factor on immune-mediated tumour control.

### Adenosine metabolism inhibition improves response to therapy and delays tumour growth

To test whether inhibiting extracellular adenosine production improves the response to ICB, we used KPAR orthotopic tumours as an immune-primed, high-RAS, RAG-4 LUAD model treated with anti-PD-1, anti-CD73, or a combination of both (**Figure 6a**). Anti-CD73 monotherapy treatment alone increased the median survival from 2.6 to 4.3 weeks (p = 0.011), with 10% (1) of mice achieving long-term survival (>9 weeks). The combination of anti-PD-1 and anti-CD73 further extended median survival to 3.9 weeks (p = 0.031), with 30% (3) of mice surviving beyond 4 weeks, suggesting additive or synergistic effects with ICB (**Figure 6b**). These findings support the hypothesis that extracellular adenosine limits the efficacy of immune checkpoint blockade in RAS-high, immune-primed tumours. We next tested whether adenosine inhibition could also enhance response to KRASG12C inhibition, which partially relies on intact anti-tumour immunity^10,11^. In KPAR^G12C^ tumours^28^, anti-CD73 improved survival in mice treated with G12Ci from 3.9 to 4.6 weeks (p = 0.002) (**Figure 6c-d**), suggesting that adenosine inhibition enhances therapeutic response even outside the ICB context.

**Figure 6.**
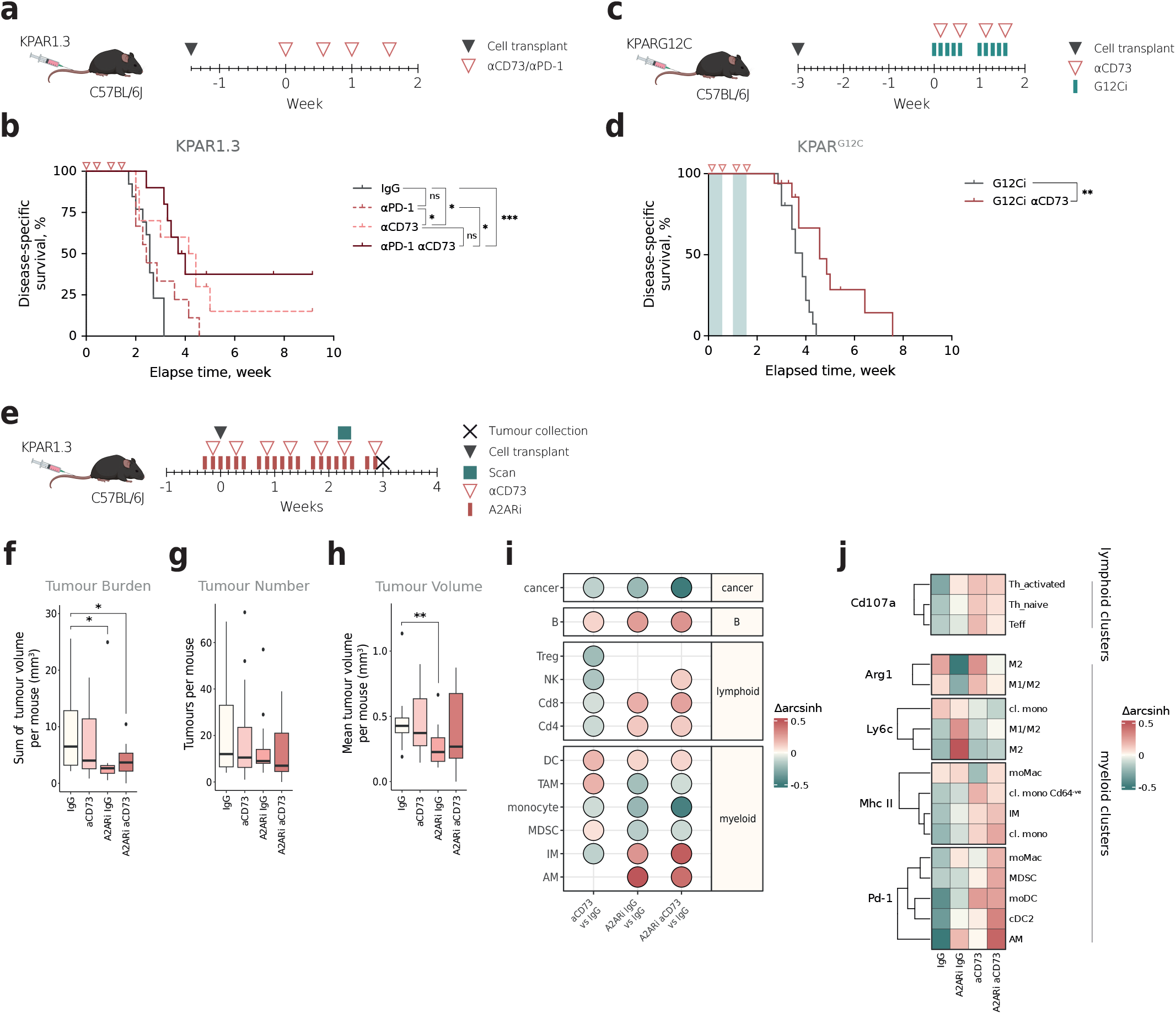
Adenosine inhibition in immune-primed KPAR mouse tumours. (a) Schematic showing the treatment and scan schedule of *in vivo* experiment. KPAR1.3 cells were transplanted 10 days before the beginning of the treatment. Mice were treated with IgG control, anti-PD-1, anti-CD73, or the combination of both anti-PD-1 and anti-CD73 at days 0, 4, 7, and 11 (indicated by pink arrowheads). (b) Kaplan-Meier plot of KPAR-tumour-bearing mice from the experiment described in (a). Mice were treated with IgG control (n = 13), anti-PD-1 (n = 9), anti-CD73 (n = 10), or the combination of both anti-PD-1 and anti-CD73 (n = 10). Statistical significance was determined by the Log-rank (Mantel-Cox) test (**p≤0.01). (c) Schematic showing the treatment and scan schedule of *in vivo* experiment. KPAR^G12C^ cells were transplanted 21 days before the beginning of the treatment. Mice received G12Ci (MRTX849) from day 0, for 2 weeks, 5 days a week (indicated by blue bars) and were treated with IgG control or anti-CD73 at days 1, 4, 8, and 11 (indicated by pink arrowheads). (d) Kaplan-Meier plot of KPAR^G12C^-tumour-bearing mice from the experiment described in (c). Mice were treated with G12Ci (indicated by light blue areas) and IgG control (n = 16) or anti-CD73 (n = 17). Combination of 2 independent experiments. Statistical significance determined by Log-rank (Mantel-Cox) test (**p≤0.01). (e) Schematic showing the treatment and scan schedule of *in vivo* experiments. Treatment started 2 days before cancer-cell transplant. Mice received anti-CD73 or IgG control twice weekly and A2ARi (AZD4635) daily, 6 days a week. (f) Boxplots with whiskers of the data collected from the experiment described in (e), showing tumour burden represented as the sum of tumour volumes per mouse. Combination of 2 independent experiments, with IgG control (n = 14), anti-CD73 (n = 14), A2ARi (n = 13), or the combination of both anti-CD73 and A2ARi (n = 15). Statistical analysis performed using a linear mixed-effects model with treatment as fixed effects, and sex and experiment as random effects. (g) Boxplots with whiskers showing tumour number represented as the sum of tumour volumes per mouse. (h) Boxplots with whiskers showing tumour number represented as the mean tumour volume per mouse. For f to h, the data were collected from the experiment described in (e). Combination of 2 independent experiments, with IgG control (n = 14), anti-CD73 (n = 14), A2ARi (n = 13), or the combination of both anti-CD73 and A2ARi (n = 15). (i) Dot plot showing the *Δ*arcsinh of pairwise comparisons in the frequency of immune cell clusters identified by spectral flow cytometry in tumours at the endpoint of the experiment described in (e). n = 8 mice per condition. Only fold changes with FDR ≤ 0.05 are shown. (j) Heatmap showing mean normalised protein expression across immune cell clusters identified by spectral flow cytometry for each experimental condition, relative to the mean expression in tumours at the endpoint of the experiment described in (e). n = 8 mice per condition. Only proteins differentially expressed across at least two conditions (FDR ≤ 0.05) are shown.

Given that RAS activation induces CD73 expression in normal pneumocytes (**Figure 4d**), we hypothesise that increasing interstitial adenosine may promote outgrowth at early stages of progression. To test this, we treated mice with anti-CD73 or the A2AR inhibitor AZD4635 prior to tumour cell implantation and monitored tumour progression by microCT (**Figure 6e**). A2AR inhibition significantly reduced tumour burden at 2.3 weeks post-transplantation, whilst anti-CD73 did not (**Figure 6f**). This effect was attributed to a reduction in tumour size, not tumour number (**Figure 6g**), indicating that A2AR inhibition impairs tumour growth rather than initial seeding.

Together, these data demonstrate that adenosine supports tumour growth during early progression and limits the efficacy of immunotherapy and KRAS inhibitor in established immune-primed tumours.

We next used high-dimensional spectral flow cytometry to profile the immune landscape after 3 weeks of adenosine metabolism inhibition (**Supp Figure 6a-f**). A2ARi reduced proportions of suppressive myeloid populations, including M2-like immunosuppressive tumour-associated macrophages (TAMs)TAMs and myeloid-derived suppressor cells (MDSCs), and increased interstitial macrophages (IMs) and alveolar macrophages (AMs) within the tumour (**Figure 6i** and **Supp Figure 6m**). Moreover, A2ARi decreased Arg-1 and increased Ly6C expressions in M2-like and hybrid M1/M2 TAMs (**Figure 6j**), indicating reduced immunosuppressive potential and a more pro-inflammatory, M1-like activation state^29,30^. By contrast, anti-CD73 increased intratumoural proportions of Monocyte-derived Dendritic cells (mo-DCs) mo-DCs, early transitioning monocytes (moMacs), cDC2s, and a subset of “pre-activated” Cd64^−^ monocytes (**Figure 6i** and **Supp Figure 6m**). Anti-CD73 also enhanced T cell degranulation, reflected by the expression of Cd107a—a marker of cytotoxic activity in lymphocytes and lysosomal mobilisation in myeloid cells—, and induced Mhc-II expression on monocytes and IM, suggesting increased antigen presentation capacity. We also observed that A2ARi increased proportions of B cells (**Figure 6i**). However, the double treatment significantly upregulated Pd-1 expression on moMacs, cDC2, and AM, potentially compromising their anti-tumour function^31^.

Notably, both treatments reduced the proportion of cancer cells, with the combination showing the most potent effect (**Figure 6i**). However, whilst the H2-Kb^lo^ cancer cluster decreased, H2-Kb^hi^ cells persisted under treatment (**Supp Figure 6h** and **m**). The H2-Kb^hi^ cluster expressed higher Cd39, Cd73, Mhc-II, Mhc-I (H2-Kb) and Pd-l1 (**Supp Figure 6c**), suggesting stronger interferon response in these cells and a greater capacity to suppress anti-tumour immunity.

These findings reveal complementary roles for anti-CD73 and A2AR inhibition in reshaping the tumour immune microenvironment, with the former activating lymphoid and anti-tumour myeloid populations, and the latter depleting suppressive myeloid cells, thereby jointly controlling tumour growth.

## Discussion

This study demonstrates that RAS transcriptional activity, quantified with the RAS84 signature, is a key determinant of the immune landscape and therapeutic response in lung adenocarcinoma (LUAD). We show that high RAS activity (RAG-3/4) is associated with T-cell infiltration, interferon responses, and improved progression-free survival following anti-PD-1 therapy. RAG-3/4 tumours that progressed upon treatment had increased interstitial adenosine (iADO) signature, which we show is driven by RAS-regulated expression of *NT5E* (CD73) and nucleoside transporters. *In vivo* experiments in preclinical models of RAG-3/4 confirm that adenosine suppresses innate and adaptive immunity and limits response to both immune checkpoint blockade (ICB) and KRAS inhibition. Together, these findings establish RAS signalling as both a driver of tumour immunogenicity and a direct promoter of immunosuppression. This duality refines our understanding of RAS biology and suggests new therapeutic avenues.

We and others have previously linked KRAS mutations to immune evasion and PD-L1 expression in cancer cells^9–12^. *KRAS* mutation was previously associated with superior survival compared with *KRAS* WT in a cohort of NSCLC patients treated with anti-PD-L1^13^; however, the data remain inconsistent across studies^14,32^. We extend these observations and demonstrate that RAS transcriptional activity, rather than KRAS mutation status, more faithfully reflects the immune phenotype and therapeutic vulnerabilities associated with oncogenic RAS signalling in LUAD. Accordingly, we observe that KRAS mutation status alone does not predict ICB benefit, whereas RAS84 stratification better separates responsive from resistant groups. The reason for that could be that many members of the RAS signalling network, other than KRAS, can be altered in lung cancer and activate RAS signalling in KRAS WT tumours^8^. By relying on transcriptional expression, our RAS classification is agnostic to genetic alterations and thus presents an advantage over KRAS mutation to study how oncogenic RAS activity shapes the immune landscape of LUAD.

Importantly, we demonstrate that increased RAS signalling correlates with an immune-primed tumour microenvironment and increased T-cell infiltration in established human LUAD and mouse lung tumours. The simultaneous immune-activating and immune-suppressive effects associated with RAS signalling reflect the complex stress response and compensatory mechanisms frequently triggered by oncogenic activation. For instance, RAS-driven proliferation induces DNA damage by exerting stress on replication forks^33^. RAS-stimulated metabolism also generates oxidative stress and reactive oxygen species (ROS)^34^. The resulting *oncogenic stress* can trigger oncogene-induced senescence (OIS) or apoptosis^35^. Yet, RAS signalling counteracts these tumour-suppressive mechanisms, notably by inducing anti-apoptotic proteins^36^, activating NF-κB^37^, and promoting epithelial-to-mesenchymal transition (EMT)—a process associated with tumour progression and escape from OIS^38^. This coordinated resistance to oncogenic stress facilitates the survival and expansion of cancer cells, leading to tumour progression. Analogously, the oncogenic stress associated with RAS activation may be sensed by the immune system, but simultaneously suppressed by direct actions of RAS signalling, such as adenosine production or PD-L1 expression

Our data also support and expand on reports that EGFR or MEK inhibition can reduce CD73 expression in lung cancer cell lines^39^. We show that oncogenic RAS signalling directly regulates adenosine-producing (CD73) and recycling (*e*.*g*., ENT family, ADA, ADK) genes in both transformed and non-transformed pneumocytes, implicating the RAS–MEK–ERK axis as a driver of iADO accumulation. Our findings place adenosine regulation as a direct output of RAS oncogenic signalling, likely occurring at early stages of tumour evolution, rather than solely a secondary consequence of immune pressure.

Our data reveal that high interstitial adenosine (iADO) accumulation may contribute to resistance to ICB in immune-primed RAG-3/4 LUAD tumours. In a preclinical model of RAG-4 tumours, blockade of adenosine production via anti-CD73 enhanced responses to both immune checkpoint blockade and KRASG12C inhibition, the latter known to require the secondary activation of the anti-tumoural immune response for maximal efficacy. These findings position adenosine as a regulator of therapeutic response in RAS-driven lung cancer.

At the cellular level, adenosine signalling suppresses both innate and adaptive immunity. Its inhibition led to enhanced recruitment of T and B cells, reduced TAMs and MDSCs, and shifted macrophage polarisation from an immunosuppressive M2-like state to a more pro-inflammatory phenotype, notably reflected by reduced Arg1 expression and increased Ly6C. Anti-CD73 treatment also activated antigen presentation on myeloid cells and decreased the proportion of M2-like immunosuppressive macrophages within tumours, consistent with a broader reprogramming of the suppressive myeloid niche.

Adenosine also contributes to tumour establishment. Oncogenic RAS activation induces the expression of adenosine-regulatory genes in normal pneumocytes, indicating that adenosine production is an early and intrinsic feature of RAS signalling.

However, inhibition of A2A receptor signalling during cancer cell seeding delayed tumour growth but did not reduce the number of tumours, indicating that adenosine promotes tumour outgrowth rather than initial engraftment and maintenance of tumour-initiating cells. Altogether, this suggests that RAS-driven adenosine production facilitates immune escape of differentiated progenitor cancer cells without being essential for cancer stem cell maintenance.

Notably, cancer cell subsets characterised by the expression of interferon proteins such as MHC class I and II, and PD-L1 persisted despite adenosine pathway inhibition. This immune-resistant population is consistent with tumour-initiating or stem-like cells, which have previously been shown to exhibit immune privilege and express immune inhibitory molecules such as PD-L1 in other cancer types^40,41^ and are associated with interferon response in NSCLC^42^. These findings suggest that while adenosine suppresses anti-tumour immunity broadly, certain highly immune-evasive subpopulations, particularly those with high IFN responses, rely on alternative immune escape mechanisms. This is consistent with data showing that loss of interferon signalling paradoxically made tumours more sensitive to T and NK cell killing through inhibition of classical and non-classical MHC class I^43^.

Together, these findings define RAS-driven adenosine accumulation as a critical driver of both tumour progression and therapy resistance, primarily through the suppression of myeloid-mediated immunity. However, subsets of cancer cells with PD-L1 upregulation persisted despite adenosine blockade in our preclinical model, suggesting that these immune-evasive populations are not dependent on adenosine for survival. Conversely, LUAD tumours resistant to ICB showed increased iADO scores, further supporting the rationale for combination therapies that integrate immune checkpoint blockade with adenosine pathway inhibition in high-RAS LUAD.

This work has direct clinical implications. First, RAS84 outperforms KRAS mutation status as a biomarker of ICB response and could help stratify LUAD patients for immunotherapy. Second, the identification of adenosine as a RAS-regulated suppressive pathway provides a rationale for combining ICB with adenosine blockade in RAS-high tumours. Early clinical trials of A2AR antagonists (e.g., ciforadenant) or anti-CD73 antibodies (e.g., oleclumab) have reported modest efficacy in NSCLC, but responses have been inconsistent^6,7^. The COAST trial (NCT03822351) reported that oleclumab combined with durvalumab (anti-PD-L1) improved PFS and objective response rates compared with durvalumab alone in unresectable stage III NSCLC patients who had not progressed after chemoradiotherapy^44^. Similarly, the combination of ciforadenant with atezolizumab, another anti-PD-L1 agent, has demonstrated encouraging tolerability and modest signs of efficacy in metastatic NSCLC^45^. Our data suggest that patient selection using RAS84 and adenosine signatures may identify the subset most likely to benefit from this combination therapy.

Third, our study highlights a therapeutic inversion: tumours with RAG-2 phenotypes— poor ICB responders—were previously among the best responders to chemotherapy. This raises the possibility of using RAS84 not only for immunotherapy stratification but also for broader treatment allocation across modalities.

Several limitations should be acknowledged. The Samsung cohort lacked complete responders, which may underestimate the predictive potential of RAS84 for durable ICB benefit. Although our orthotopic transplant models better capture the immunogenicity observed in LUAD, they do not model early tumour evolution or immune surveillance at initiation. Genetically engineered mouse models, whilst less immunogenic, may still be required to fully explore how RAS-driven adenosine regulation shapes the earliest phases of tumour development. Finally, although we demonstrate consistent transcriptional and functional effects across multiple systems, prospective clinical validation of RAS84 in immunotherapy trials is essential.

The predictive value of RAS84 in the context of therapy should be tested in prospective LUAD cohorts. Integrating RAS84 with other immune signatures, Tumour Mutation Burden (TMB) or adenosine scores could refine patient stratification for combined ICB and adenosine-targeting therapies. Future studies should also focus on dissecting the temporal dynamics of RAS-induced immunogenicity and immune suppression, particularly during tumour initiation when immune escape mechanisms are initially established in response to oncogenic RAS activation. Gaining new insights into how RAS shapes early tumour evolution and establishes the immune landscape could help identify therapeutic windows for early intervention and guide the rational design of combination therapies targeting both tumour-intrinsic and immune-mediated resistance mechanisms.

In conclusion, this work establishes oncogenic RAS signalling as a key determinant of tumour immunity and positions RAS transcriptional activity as a surrogate biomarker to guide immunotherapy and uncover novel mechanisms of RAS-driven immune evasion.

## Methods

### Samsung Cohort Analysis

#### Patients

314 Advanced NSCLC treated with CPI (PD-L1 or programmed cell death protein 1 [PD-1] monotherapy, or in combination with cytotoxic T-lymphocyte antigen 4 inhibitor) at the Samsung Medical Center and Seoul National University Hospital between June 2014 and February 2020 were included. The medical records were retrospectively reviewed. Tumour response was assessed using the Response Evaluation Criteria in Solid Tumours version 1.1. This study was approved by the Institutional Review Board of the Samsung Medical Center (IRB no. SMC-2018-03-130), and written informed consent was obtained from all enrolled patients.

#### Survival analysis

Progression-free survival (PFS) was defined as the duration from the start of CPI until progression. Kaplan–Meier estimates and log-rank tests were used for survival analysis. The Cox proportional hazards regression model was used to calculate the hazard ratio (HR). Statistical significance was set at *P* < 0.05. R version 4.0.2 was used for statistical analyses.

#### Whole-transcriptome sequencing (WTS) and data analysis

RNA was purified from formalin-fixed paraffin-embedded (FFPE) or fresh tumour samples using the AllPrep DNA/RNA Mini Kit (Qiagen, USA). RNA concentration and purity were measured using a NanoDrop and Bioanalyzer (Agilent, USA). The library was prepared following the manufacturer’s instructions using either the TruSeq RNA Library Prep Kit v2 (Illumina, USA) or the TruSeq RNA Access Library Prep Kit (Illumina, USA). The reads from FASTQ files were mapped against hg19 using the 2-pass mode of STAR version 2.4.0. The first pass aligned reads to the hg19 genome reference and generated the sample-specific reference using the hg19 genome. The second pass was used to align the reads to the newly generated hg19 genome and the sample-specific reference. RNA-SeQC was conducted to control the quality of the BAM files. Raw read counts mapped to genes were analysed for transcript abundance using RSEM version 1.2.18, and poorly expressed samples were eliminated when the read count criterion was < 1M. ComBat was used to merge gene expression generated with different platforms to eliminate batch effects between different datasets.

### Bioinformatics and Computational Analyses

#### RAS84 and mRAS79 Classification

RAS activity was stratified into RAS-activity groups (RAG-0 to RAG-4) using the RAS84 gene signature, following hierarchical clustering with Euclidean distance and ward.D2 agglomeration (hclust function, R). RI (RAS Index) values were calculated as the mean expression of RAS84 genes for each sample, as described previously^15^. Gene expression deviations and tumour purity corrections were performed using DESeq2 (v1.24.0).

#### Differential gene expression (DGE) analysis and signature score calculation

To identify DGE between the responders and the non-responder we used the DESeq2 package with default option. Significant genes were tested using the two-sample t-test in each group. p value < □0.05 was considered significant.

#### Gene Set Enrichment Analysis (GSEA)

Hallmark pathway analysis was conducted using GSEA with MSigDB gene sets. Enrichment scores and FDR were calculated to assess differences across RAGs and treatment groups.

#### HLA LOH analysis

HLA LOH calls for HLA A, B and C genes for the TRACERx421 cohort were taken from a previous TRACERx study^46^, generated using the MHC Hammer pipeline. Only tumour samples for which (i) at least one gene of HLA A, B or C had passed whole-exome sequencing, and (ii) for which a RAS84 group had been assigned were included in our analysis. Sample-level HLA LOH calls were then calculated by assigning a sample as having HLA LOH if any one or more of the HLA A, B or C genes for which HLA LOH data were available for the sample showed evidence of HLA LOH. To assess whether an association existed between sample-level RAS84 status and HLA LOH status, a generalised linear mixed-effects model was applied (*glmer* R function, *lme4* R package; family = binomial, control = glmerControl(optimizer = “bobyqa”), nAGQ = 10)), comparing the frequency of HLA LOH in each RAS84 group with all other RAS84 groups combined. Tumour ID was included in the model as a random effect to account for varying numbers of tumour regions per patient. Each full model was then compared to a null model in which RAS84 group was not included as an independent variable, and the ANOVA p-value comparing full to null model used to quantify significance (** p<0.01, * p<0.05).

#### CCLE protein expression data

CCLE protein expression data were downloaded from https://gygi.hms.harvard.edu/publications/ccle.html. We used the normalised protein quantification of lung cell lines from Nusinow *et al*.^47^. We calculated the Pearson correlation between CD73 protein expression and *NT5E* mRNA expression across all cell lines for which CCLE RNA-Seq and protein expression were available (n = 63 NSCLC cell lines and n = 38 lung adenocarcinoma cell lines).

#### Adenosine score calculation

We measured the signature score in each sample using the VST value from the DESeq2 R packages. We generated Z-score matrix from the VST value matrix and calculated the mean value of the Adenosine signature gene expression.

#### Spectral-flow data analysis

After pre-gating on live cells in FlowJo, we created a flowSet object from the FCS files using flowCore (2.16.0). We used flowVS (1.36.0)^48^ to estimate channel-specific arcsinh cofactors and applied these to transform the intensity values. We applied a manually derived cofactor in cases where flowVS was unable to estimate a cofactor that centred the background signal around zero (**Supp. Table 1**). We transformed the flowframe objects to SingleCellExperiment (1.34.0) objects and subsampled 5,000 cells from each sample. We merged the samples. We clustered the cells using Louvian clustering (Rphenograph 0.99.1)^49^ across a set of lineage-specific markers (**Supp. Table 2**) to identify broad immune-cell types. We annotated the clusters by lineage marker expression and separated the cells into their lineage groups (AM, B, Cancer, IM, MonMacDC, MDSC, TNK and Other) (**Supp. Table 3**). We reclustered the lineage groups using lineage-specific markers (**Supp. Table 4**) and annotated the clusters (**Supp. Table 5**). We identified treatment-dependent changes in cell-type population sizes by fitting a Poisson generalised linear model to the cell counts, counts ~ treatment and using the total number of cells as the offset. We applied multiple testing correction and selected significant cell types using a < 0.05 FDR filter. We carried out this analysis within each lineage group and across all cells. We identified differential antibody expression across the four treatment conditions. We calculated the mean antibody intensity values for each cell type within each sample. We tested for changes in expression by fitting a linear model to the mean intensity values across treatments. We applied multiple testing and selected antibodies using a < 0.05 FDR value. We carried out the analysis in R-4.4.1^50^.

### Cell Culture and In Vitro Studies

#### Cell Culture and RAS Inhibition

KPAR1.3 and KPARG12C cells ^28^ were maintained in DMEM supplemented with 5% FBS and Glutamax. H23, H1792, H358, and SW873 cells were obtained from the Francis Crick Institute Cell Services. The cells were maintained in RPMI 10% FBS and treated with ARS853, ARS1620, or MRTX849 at the indicated concentration for the indicated time.

#### In Vitro Treatment with MRTX1257 and AMP Metabolism Assay

H1792 cells were seeded in four replicates at a density of 80,000 cells per well in 12-well plates, cultured in RPMI medium supplemented with 5% FBS and Glutamax, and incubated for 24 hours. Cells were treated with 200 nM MRTX1257 or vehicle control (DMSO, final concentration 0.004%). Medium was replaced 24 hours post-seeding before drug treatment, and additional doses of MRTX1257 were added every 24 hours without changing the medium. At 72 hours post-treatment initiation, the medium was removed, and cells were washed twice with PBS. Fresh RPMI medium containing 10 µM AMP was added to each well. Supernatants were collected at 5, 10, 30, and 60 minutes, centrifuged at 500 g, 4°C, for 5 minutes, and immediately frozen at −80°C for LC-MS analysis. Cell counts from three additional wells per condition were performed to normalise AMP metabolism data.

### Biochemical and Molecular Analyses

#### Western Blot

Protein lysates were prepared using RIPA buffer supplemented with protease and phosphatase inhibitors (Sigma). Proteins were resolved on 4-12% Bis-Tris gels (Thermo Fisher) and transferred to PVDF membranes. Primary antibodies targeting CD73, pERK1/2, and vinculin (loading control) were visualised using HRP-conjugated secondary antibodies on a Chemidoc imaging system (Bio-Rad).

#### qPCR for Gene Expression Analysis

RNA was extracted using the RNeasy kit (Qiagen), and cDNA synthesis was performed using Maxima First Strand cDNA Synthesis Kit (Thermo Fisher). qPCR was carried out using SYBR Green Master Mix (Thermo Fisher) on a StepOnePlus system (Applied Biosystems). Data were normalised to housekeeping genes *GAPDH* and *ACTB*. The primers used in this study are referenced in **Supp. Table 6**.

#### Analysis of Adenosine and AMP by Liquid Chromatography-Mass Spectrometry (LC-MS)

To 50 µL of collected media, 150 µL of ice-cold methanol was added for protein precipitation, followed by thorough vortexing. Biphasic partitioning was performed by adding 100 μl of water followed by 50 μl of chloroform. Tubes were vortexed and centrifuged at 10,000 ×g (10 min, 4oC) and the upper polar phase was transferred to glass vials with inserts. The LC-MS method was adapted from Fets and colleagues ^51^ . Briefly, samples were injected into a Dionex UltiMate LC system (Thermo Scientific) using a ZIC-pHILIC (150□mm□× □4.6□mm, 5-μm particle) column (Merck Sequant). A 15-min elution gradient was used (80% solvent A to 20% solvent B), followed by a 5-min wash (95:5 solvent A to solvent B) and 5-min re-equilibration; solvent A was 20□mM ammonium carbonate in water (Optima HPLC grade, Sigma Aldrich) and solvent B was acetonitrile (Optima HPLC grade, Sigma Aldrich). Flow rate was 300□µl/min; column temperature was 25□°C; injection volume was 10□µl; and autosampler temperature, 4□°C. MS was performed in positive mode using a TSQ-Quantiva (Thermo Scientific), parameters were as follows: spray voltage 3500 V; capillary temperature 375 °C; vaporizer temperature 275 °C; sheath, auxiliary and sweep gases were 45, 16 and 5 arbitrary units, respectively; CID gas 1.5 mTorr, Q1 and Q3 resolution (FWHM) of 0.7 and 1.2, respectively. Analysis was performed in selected reaction monitoring (SRM) mode using the mass transitions for adenosine (precursor 268 m/z and product ions 119 and 136 m/z) and AMP (precursor 348 m/z and product 136 m/z). Collision energies were manually optimized using commercial standards and dwell times were set to 40 ms for each transition. Data was acquired using Xcalibur 4.0.27.10 software (Thermo Scientific). Quality control samples were prepared by pooling small aliquots of all samples analysed to ensure analytical accuracy, and the concentration of the compounds was determined using calibration curves generated from media spiked with commercial standards. Qualitative and quantitative analysis was performed using Free Style 1.8 and TraceFinder 5.1 software (Thermo Scientific), according to the manufacturer’s workflows.

#### Spectral flow cytometry

Mice were euthanised using Schedule 1 methods. Lung tumours were dissected, and all tumours from one lung were pooled. Tumours were finely minced and digested in HBSS containing collagenase (1□mg/mL; Thermo Fisher Scientific) and DNase I (50□U/mL; Life Technologies) for 45□minutes at 37□°C. The resulting cell suspension was filtered through 70□μm strainers (Falcon), and red blood cells were lysed using ACK buffer (Life Technologies). Cells were incubated with anti-CD16/32 antibody (BioLegend) for 5□minutes at 4□°C to block Fc receptors, followed by staining at room temperature with surface antibodies and Zombie NIR fixable viability dye (BioLegend). After staining, samples were fixed in Fix/Lyse solution (eBioscience). For intracellular staining, cells were fixed and permeabilised using the Foxp3/Transcription Factor Fixation/Permeabilisation Kit (Invitrogen), following the manufacturer’s protocol, before incubation with intracellular antibodies. Single-stain compensation controls were prepared using mouse spleen cells or OneComp eBeads (Invitrogen). Antibodies used are listed in (**Supp. Tables 7**). Samples were resuspended in FACS buffer and analysed using a 5-laser Aurora spectral cytometer (Cytek).

### In Vivo Experiments

#### Orthotopic lung cancer mouse models and tumour imaging

1.5×10^5 KPAR1.3 or KPARG12C cells in 200 μl phosphate-buffered saline (PBS) were orthotopically transplanted into syngeneic C57BL/6J mice via tail vein injection. Tumour burden was monitored using Quantum GX2 micro-CT imaging (Perkin Elmer) as previously described^52^ and through weighing twice weekly. Tumour volume was reconstructed and analysed using Analyse software (AnalyzeDirect).

All animal studies adhered to the UK Home Office and institutional ethical guidelines under a UK Home Office-approved project license and approved by the Biological Research Facility at the Francis Crick Institute. Experiments were carried out using 8–10-week C57BL/6J mice. Mice housing is maintained with a 12–12 h light-dark cycle and in specific-pathogen-free conditions, and each cage contains nesting and is individually ventilated and never exceeds five mice. Humidity and temperature are maintained according to UK Home Office guidelines, 20–24 °C and 45–65%, respectively.

#### Drug Treatments and Inhibitor Studies in Mice

Mice received 50 mg/kg of MRTX849 (MedChemExpress) prepared in 10% captisol diluted in 50 mM citrate buffer pH 5.0 by oral gavage daily, 5 days a week; AZD4635 (provided by AstraZeneca under a collaboration research agreement) prepared in drinking water by oral gavage daily 6 times a week. For treatments with antibodies, mice received 200 μg anti-PD-1 (clone RMP1-14, cat. BE0146, BioXcell) and/or 250 μg anti-CD73 (clone 2C5-IgG1, provided by AstraZeneca under a collaboration research agreement) or the corresponding IgG controls (rat IgG2a and mouse IgG1, respectively). Antibodies were dissolved in PBS and administered via intraperitoneal injection (100 μl) twice weekly.

### Statistical Analyses

#### Kaplan-Meier Survival Analysis

We monitored disease-specific mouse survival, indicated by a 15% loss of body weight and associated with lung tumour burden, validated by microCT scans or post-mortem analysis. Disease-specific survival from *in vivo* experiments was estimated using Kaplan-Meier analysis. Hazard ratios were calculated using Cox proportional hazards modelling in GraphPad Prism v9. Log-rank tests were used for group comparisons.

#### General Statistical Analyses

All statistical tests for *in vitro* and *in vivo* analyses, including ANOVA, t-tests, and chi-squared tests, were performed in R (v4.2.0) or Prism (v9). P-values were adjusted for multiple comparisons using the Benjamini-Hochberg method. Data are expressed as mean ± SEM unless otherwise specified.

## Data and code availability

KPAR RNA-Seq data samples are available at GEO accession GSE199582. The five previously unpublished KPAR samples (MOL38-42) are available at GEO accession GSE310418. 3LL samples are available from GSE183548.

Custom code used in this study is available at our GitHub repository [https://github.com/phileastbioinf/decarne-et-al-RAS84-immune].

## Acknowledgements

We thank the core facilities at the Francis Crick Institute, including the Biological Research Facility, Scientific Computing, Flow Cytometry, Experimental Histopathology, Cell Science, and In Vivo Imaging Science Technology Platforms (STP). We acknowledge the Genomics STP, and particularly Deb Jackson, Daniel Leonce and Marg Crawford, for their contributions to mRNA library preparation and sequencing. We acknowledge the Bioinformatics and Biostatistics STP for their analysis support. We thank the members of the Oncogene Biology Laboratory for their discussions and critical reading of the manuscript.

## Author contributions

S.C.T., P.E. and J.D. conceived the study. S.C.T. and J.D. supervised the work and acquired funding. S.C.T. performed experiments and computational analyses. C.E.P., M.S.d.S., H.C., E.C., S.R., C.M., S.L., D.C., J.B., M.T., R.B., B.T. and M.S. performed experiments. P.E., T.G., C.E. and E.C. carried out computational and statistical analyses. J.E. provided key reagents, and S.-H.L. and C.S. provided patient cohort data. S.C.T., P.E., C.E.P., M.S.d.S., H.C., E.C., T.G., B.T., M.M.-A. and J.D. drafted or revised the manuscript. All authors reviewed and approved the final version.

