## Supplemental Figure 1-6 and legends for "Oncogenic RAS activity is linked to immune priming and adenosine-driven immune evasion in lung adenocarcinoma"

**Supplementary Figure 1** | Related to figure 1

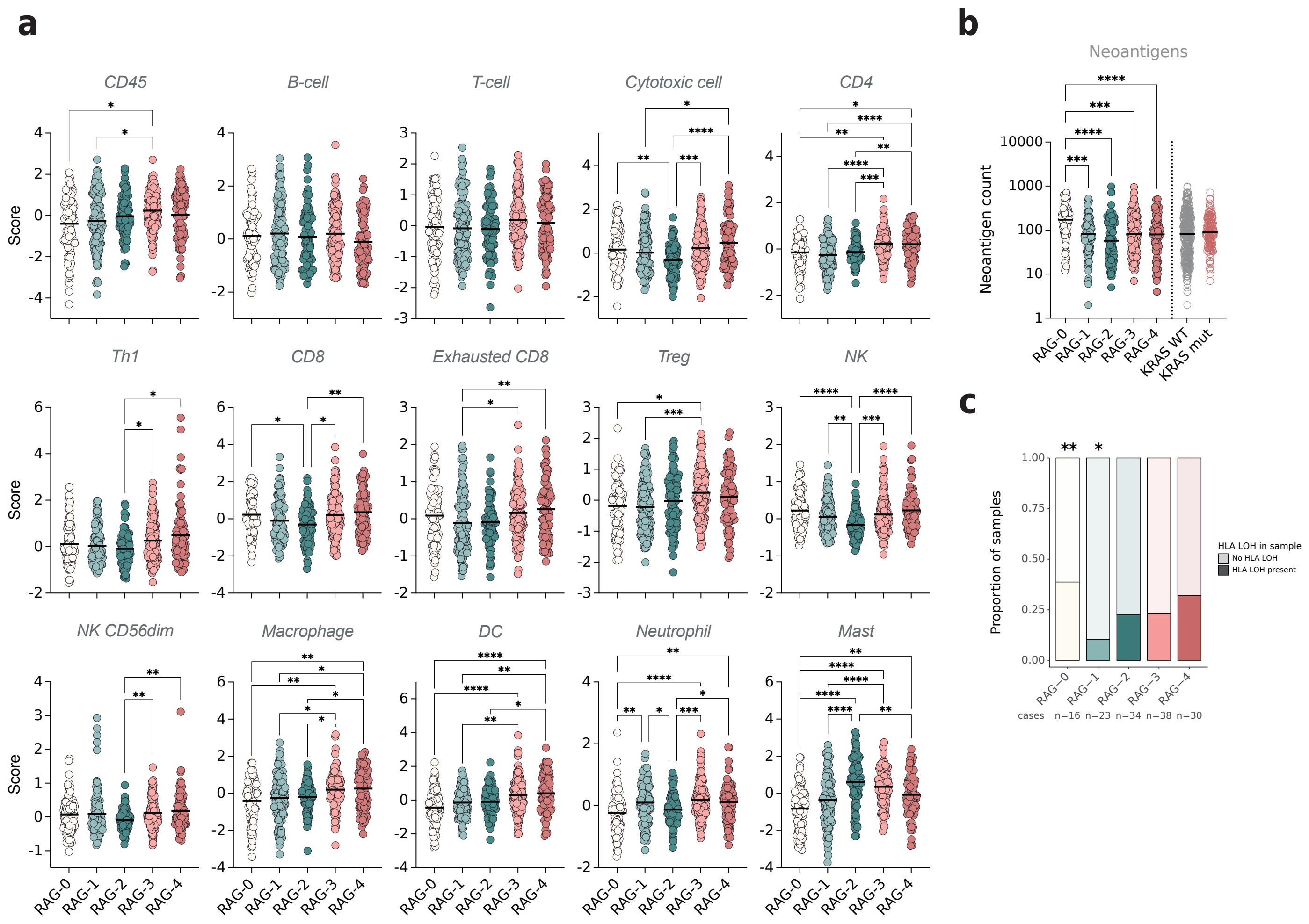

### Supplementary Figure Legends

**Supplementary figure 1 | Related to figure 1**

**(a)** Danaher and Davoli immune-cell scores scaled to median gene values across samples, individual tumour scores, and mean per RAG in LUAD tumours (TCGA). Non-parametric Kruskal-Wallis test, Dunn's multiple comparison test (*$p \leq 0.05$, **$p \leq 0.01$, ***$p \leq 0.001$, ****$p \leq 0.0001$).

**(b)** Neoantigen count from Thorsson et al. (2018)[18], median and individual count per RAG or KRAS mut and WT groups in LUAD tumours (TCGA). Non-parametric Kruskal-Wallis test, Dunn's multiple comparison test (***$p \leq 0.001$, ****$p \leq 0.0001$).

**(c)** Association between HLA loss of heterozygosity (LOH) and RAS activity groups (RAGs) in LUAD (TRACERx). Frequency of tumour samples with HLA LOH on at least one of the three HLA-A, -B, and -C genes stratified according to their RAG classification (RAS84). Generalised linear mixed-effects model comparing each RAG group against all other RAGs combined, accounting for tumour regions per patient as a random effect, ANOVA comparison with a null model (**$p < 0.01$, *$p < 0.05$).

**Supplementary Figure 2** | Related to figure 2

**a**

| **Characteristics** | **RAG-0** ***63 (20%)*** | **RAG-1** ***65 (21%)*** | **RAG-2** ***47 (15%)*** | **RAG-3** ***95 (30%)*** | **RAG-4** ***44 (14%)*** |
|---|---|---|---|---|---|
| Age | 60 (29-81) | 62 (34-84) | 57 (39-72) | 60 (34-79) | 61 (30-84) |
| Sex, *n* (%) | | | | | |
| Male | 47 (75%) | 45 (69%) | 23 (49%) | 64 (67%) | 30 (68%) |
| Female | 16 (25%) | 20 (31%) | 24 (51%) | 31 (33%) | 14 (32%) |
| Smoking status | | | | | |
| Never | 16 (25%) | 23 (35%) | 26 (55%) | 31 (33%) | 13 (30%) |
| Ever | 47 (75%) | 42 (65%) | 21 (45%) | 64 (67%) | 31 (70%) |
| Treatment line | | | | | |
| 1 | 22 (35%) | 31 (48%) | 17 (36%) | 48 (51%) | 25 (57%) |
| 2 | 20 (32%) | 13 (20%) | 8 (17%) | 20 (21%) | 4 (9%) |
| 3 | 21 (33%) | 21 (32%) | 22 (47%) | 27 (28%) | 15 (34%) |
| Driver mutation | | | | | |
| EGFR | 19 (30%) | 9 (14%) | 21 (45%) | 22 (23%) | 7 (16%) |
| KRAS | 2 (3%) | 8 (12%) | 2 (4%) | 13 (14%) | 11 (25%) |
| Others | 6 (10%) | 3 (5%) | 11 (23%) | 14 (15%) | 7 (16%) |
| Wild type | 36 (57%) | 45 (69%) | 13 (28%) | 46 (48%) | 19 (43%) |
| Response | | | | | |
| PR | 15 (24%) | 15 (23%) | 1 (2%) | 39 (41%) | 18 (41%) |
| SD | 10 (16%) | 12 (18%) | 13 (28%) | 25 (26%) | 7 (16%) |
| PD | 38 (60%) | 38 (58%) | 33 (70%) | 31 (33%) | 19 (43%) |

**b**

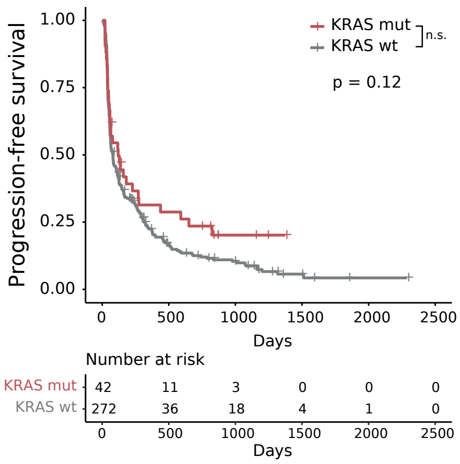


**c**

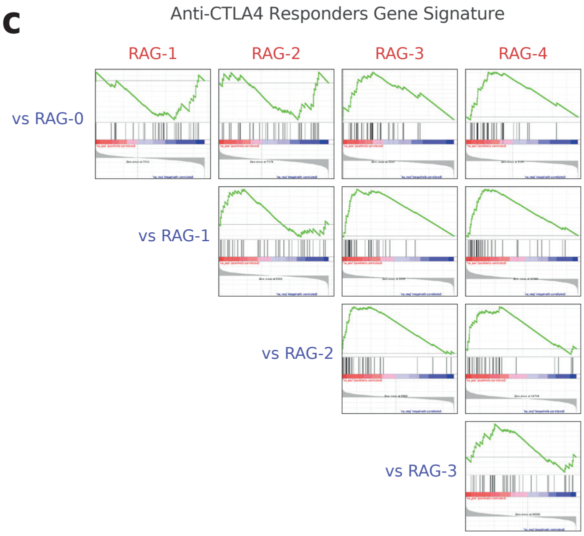

Supplementary figure 2 | Related to figure 2

**(a)** Samsung patient demographics and clinical characteristics stratified using RAS84 (RAG-0 to RAG-4). Data are presented as the number of patients with percentages in parentheses, where applicable. Age is reported as median (range). The characteristics include sex distribution, smoking status, response to anti-PD-1 therapy (as measured by progressive disease [PD], stable disease [SD], partial response [PR]), the presence of other mutations such as EGFR and KRAS, and the line of treatment received. This cohort includes n = 314 patients.

**(b)** GSEA enrichment plots from GSEA of anti-CTLA-4-responders gene signature (Ock et al. 2017)[23] in LUAD (TCGA) RAGs.

**(c)** GSEA enrichment plots from GSEA of anti-CTLA-4-responders gene signature (Ock et al. 2017)[23] in LUAD (TCGA) KRAS mutants.

**Supplementary Figure 3** | Related to figure 3

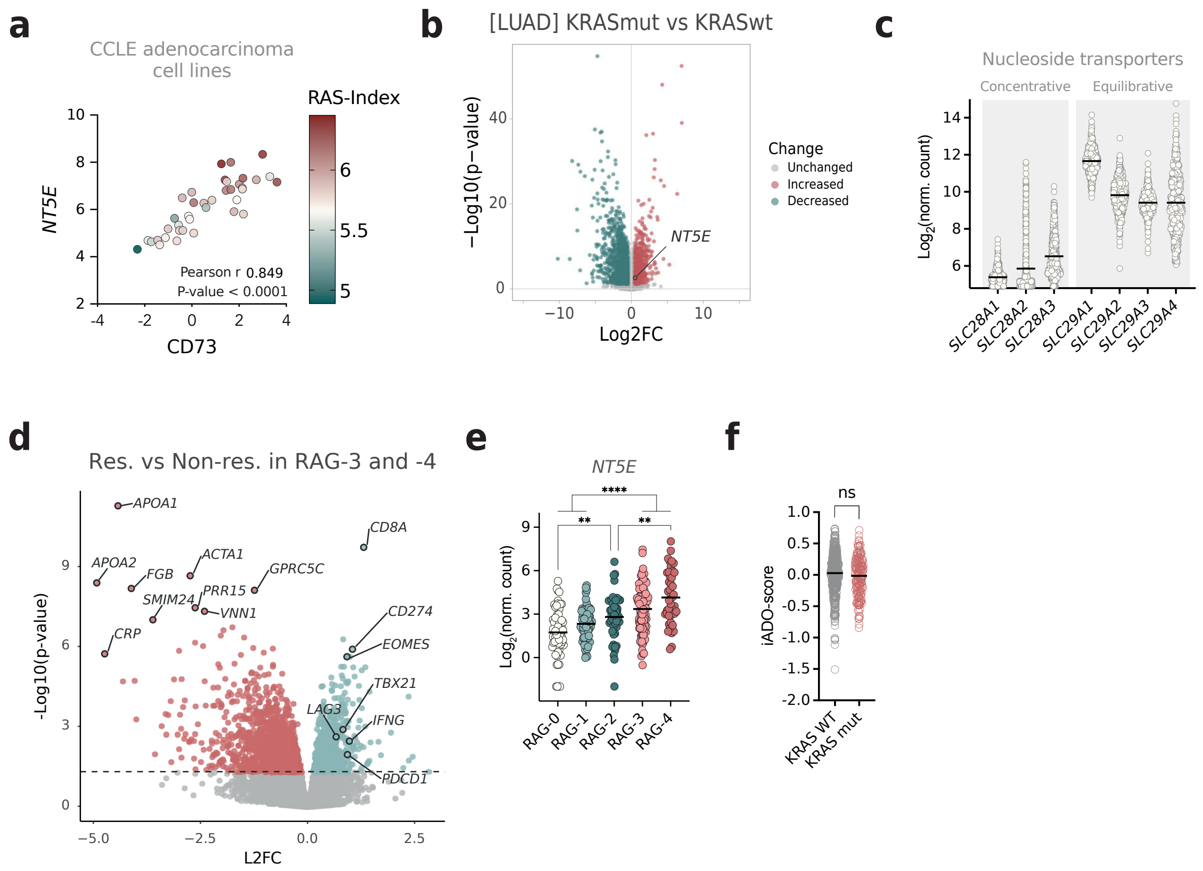

Supplementary figure 3 | Related to figure 3

**(a)** Scatterplot showing relative expressions of *NT5E* mRNA (y-axis), its protein CD73 (x-axis), and the RAS-index (colour scale) of all CCLE adenocarcinoma cell lines with both data available (n = 38).

**(b)** Volcano plot showing Differential Gene Expression between KRASmut and KRASwt in LUAD TCGA, Log2FC and -log10(p-value). Unchanged genes in grey are genes with a log2FC <0.3 or a -log10(p-value) <1.3, equivalent to a p-value >0.05.

**(c)** Concentrative and equilibrative nucleoside transporter gene expression presented as mean per group (cohort) and individual tumour values in LUAD (TCGA) (n = 502).

**(d)** Volcano plot of the gene differential expression between responders [PR] (n = 57) and non-responders [SD]+[PD] (n = 82) belonging to RAG-3 and RAG-4. Red genes are upregulated in the responders, teal genes are upregulated in non-responders, and grey genes are unchanged with a p-value < 0.05.

**(e)** *NT5E* mean and individual expression in pre-treatment tumour biopsy per RAG in LUAD tumours (Samsung). Brown-Forsythe and Welch ANOVA tests corrected for multiple comparisons using Games-Howell's tests (**$p \leq 0.01$, ***$p \leq 0.001$, ****$p \leq 0.0001$).

**(f)** Interstitial adenosine signature score (iADO-score) (Sidders et al. 2019)[25] mean and individual expression in KRAS mutant and KRAS WT LUAD tumours (TCGA). Brown-Forsythe and Welch ANOVA tests corrected for multiple comparisons using Games-Howell's tests (**ns** $p \geq 0.05$).

**Supplementary Figure 4** | Related to figure 4

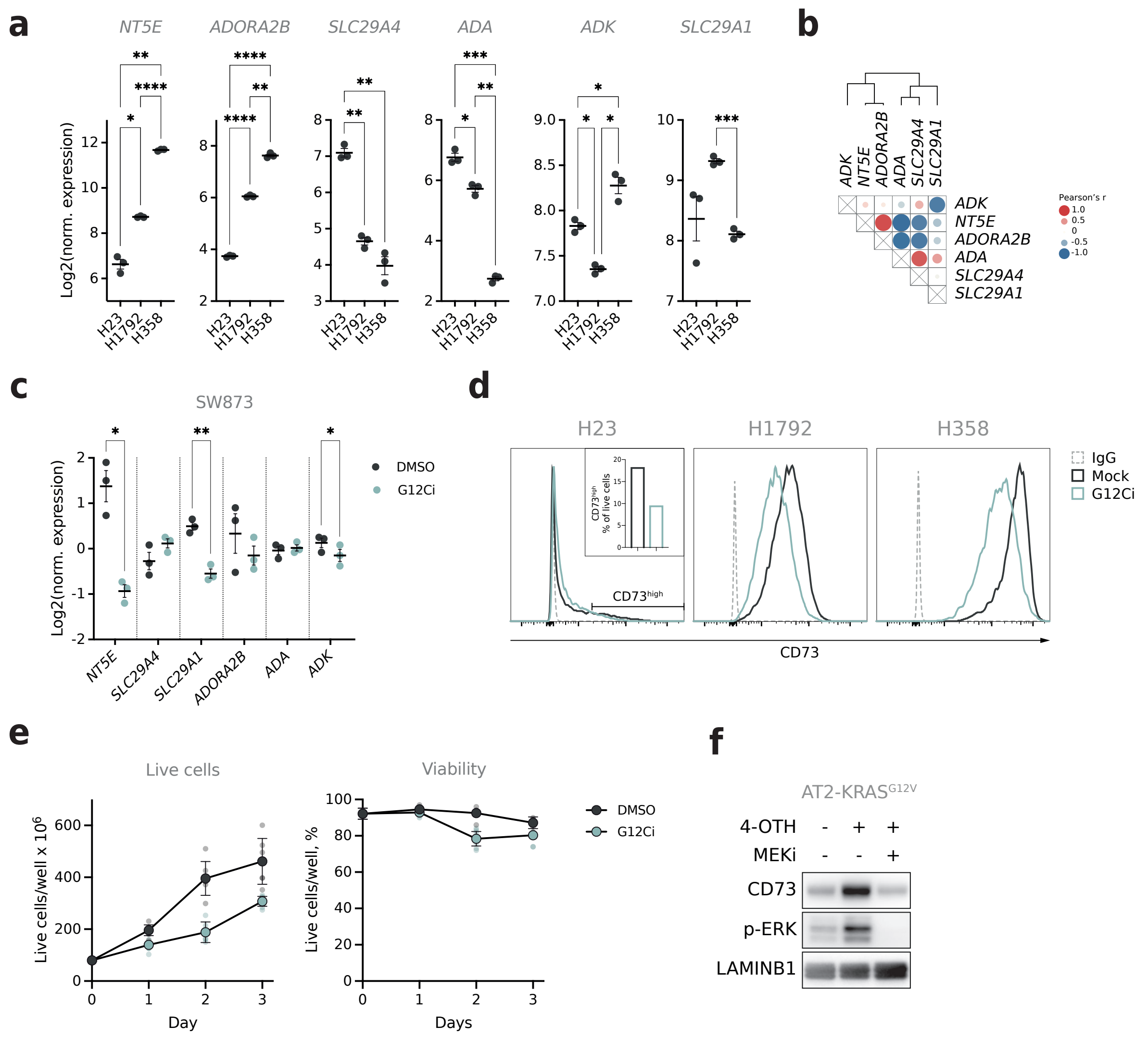

Supplementary figure 4 | Related to figure 4

**(a)** Adenosine metabolism-related gene basal expression in indicated lung cancer cell lines from qPCR. One-way ANOVA, with the Geisser-Greenhouse correction. Tukey's multiple comparisons test (*****$p \le 0.05$, ******$p \le 0.01$, *******$p \le 0.001$, ********$p \le 0.0001$).

**(b)** Heatmap showing Pearson correlations calculated using z-score normalised expression from qPCR data across H23, H1792 and H358 treated or not with 10 µM ARS853 for 24h.

**(c)** Adenosine metabolism-related gene expression in colorectal cancer cell line SW873 from qPCR, mean ±SEM per treatment group, dots represent independent biological replicates (n = 3). The cell line received DMSO or 10 µM ARS853 for 24h. Two-sided p-value with unequal variances *t*-test performed between treatment groups (*****$p \le 0.05$, ******$p \le 0.01$).

**(d)** CD73 membrane expression measured by flow cytometry in the indicated cell lines. The cell lines received DMSO or 2.5 µM ARS1620 for 72h.

**(e)** Count of live cells and viability over time. H1792 cells pre-treated with DMSO or 200 nM MRTX1257 for 72h. Mean ±SEM of 3 biological replicates (large dots) and 2 technical replicates (small dots) per time and treatment group.

**(f)** Western blot showing CD73, p-ERK1/2 (Thr202/Tyr204) and LAMINB1 in AT2-KRASG12V-ER cell line. $KRAS^{G12V}$ was induced with 100 nM 4-OTH for 24h in the presence or not of 10 nM trametinib.

**(g)** *Cd8b1*, *Gzma*, and *Pfr1* expression from qPCR mean ±SEM per treatment group. Large dots represent the mean per mouse (n = 3 per group), and small dots represent individual tumour expression. KP-tumour-bearing mice received 4 doses of trametinib over 3 days. Brown-Forsythe and Welch ANOVA tests performed on mean per mouse, corrected for multiple comparisons using Games-Howell's tests (**ns** $p > 0.05$).

**(h)** *Cd3* and *Ncr1* expression from qPCR mean ±SD per treatment group. Large dots represent the mean per mouse (n = 3 per group), and small dots represent individual tumour expression. KP-tumour-bearing mice received 4 doses of trametinib over 3 days. Brown-Forsythe and Welch ANOVA tests performed on mean per mouse, corrected for multiple comparisons using Games-Howell's tests (**ns** $p > 0.05$, *****$p \le 0.05$, ****** $p \le 0.01$).

#### **Supplementary Figure 5** | Related to figure 5

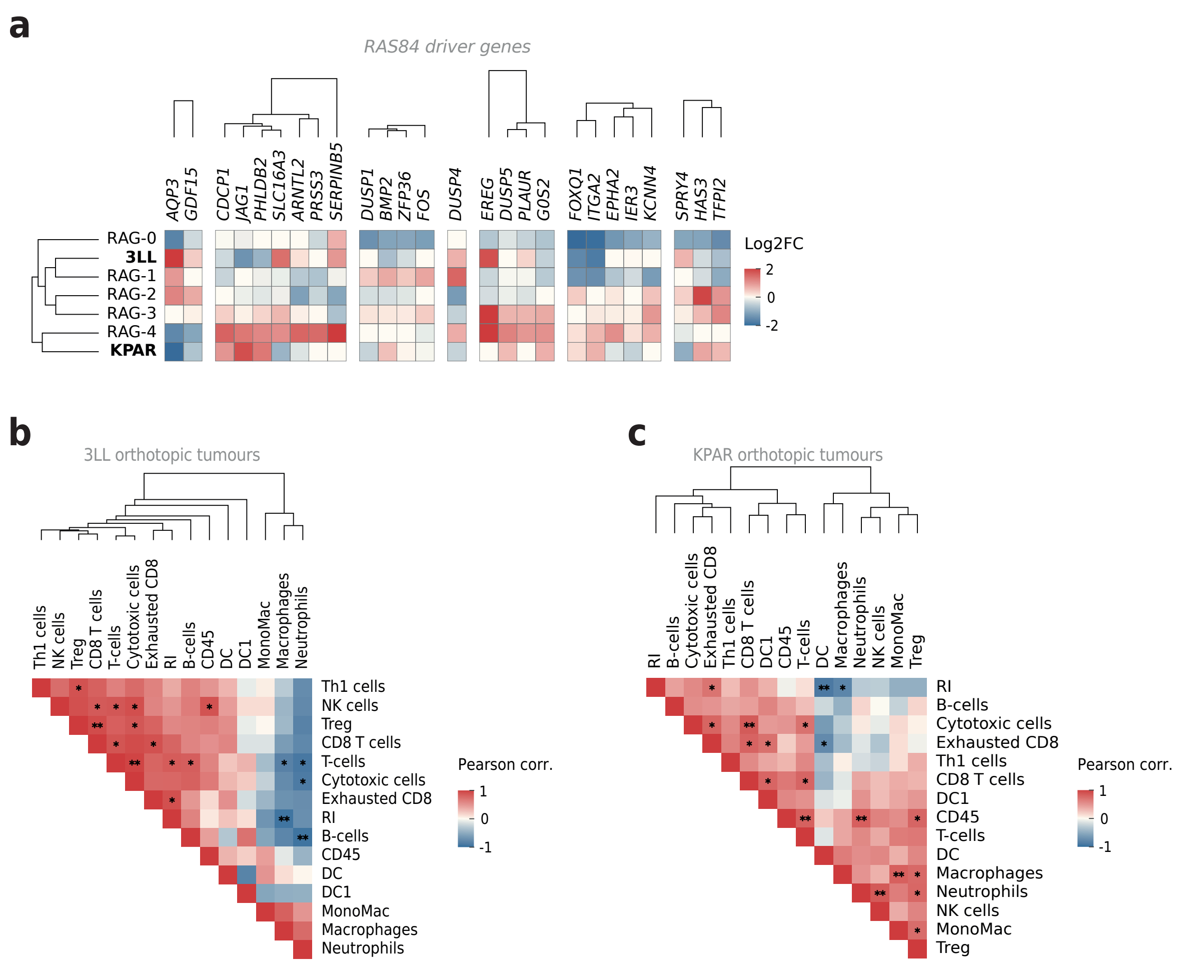

Supplementary figure 5 | Related to figure 5

**(a)** Heatmap showing variant mean RAS84 gene expression clusters across human RAGs and mouse KPAR and 3LL models with Euclidian unsupervised clustering.

**(b)** Heatmap showing the Pearson correlation between RAS-index (RI) and immune cell scores (adapted from Danaher et al. 2017)[19] across 3LL tumours (n = 6 from 3 mice).

**(c)** Heatmap showing the Pearson correlation between RAS-index (RI) and immune cell scores (adapted from Danaher et al. 2017)[19] across KPAR tumours (n = 9 from 3 mice).

**Supplementary Figure 6** | Related to figure 6

**a**

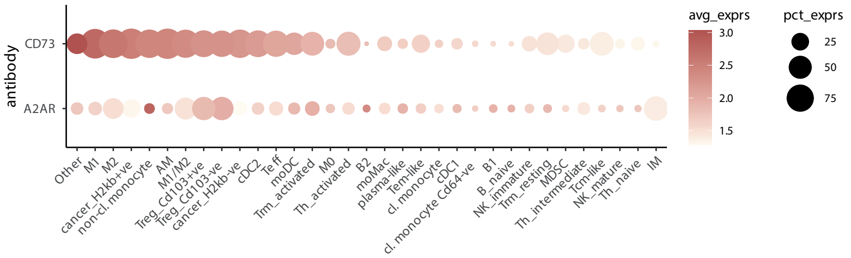


**b**

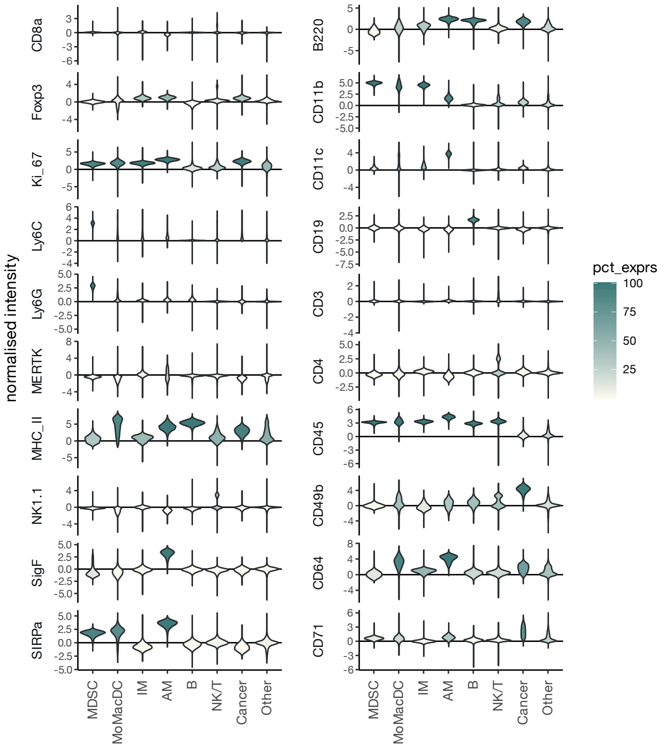

**Supplementary Figure 6** | Related to figure 6

c

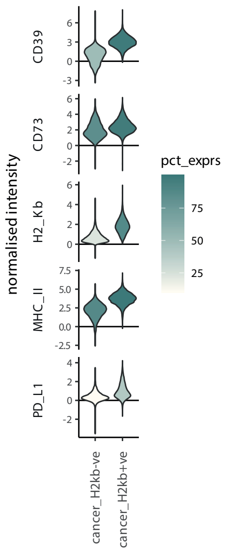


d

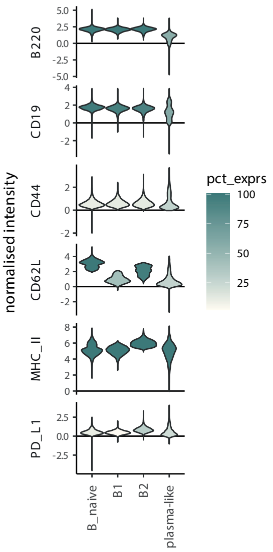


e

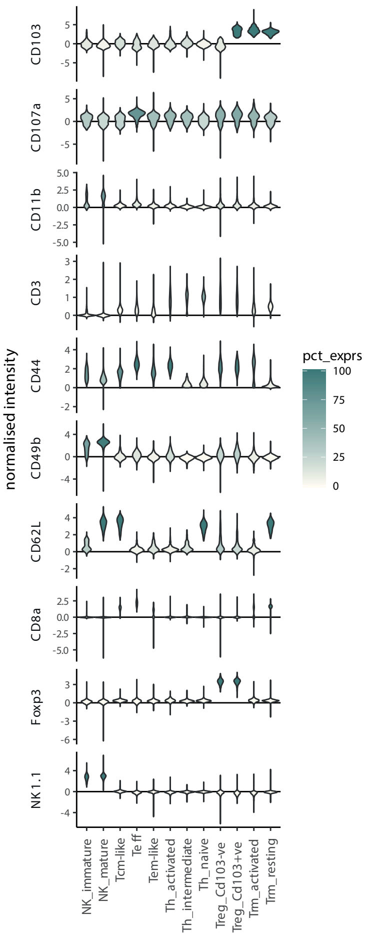


f

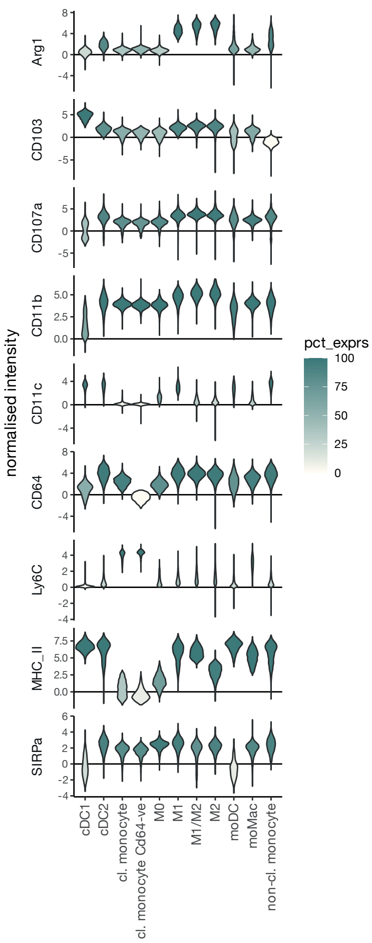

**Supplementary Figure 6** | Related to figure 6

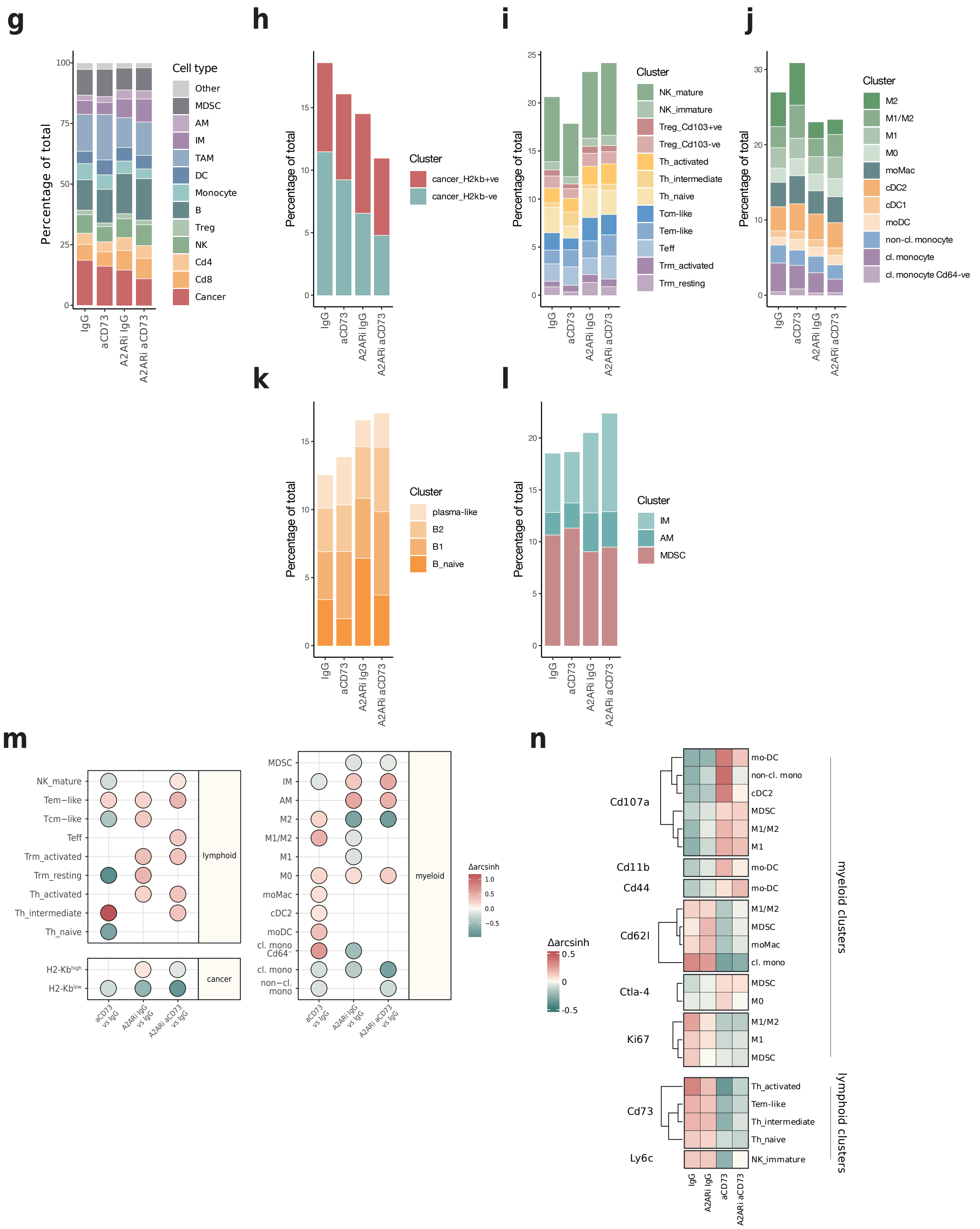

**Supplementary figure 6 | Related to figure 6**

**(a)** Bubble plot showing CD73 and A2AR expression across all cell clusters identified by spectral flow cytometry in tumours from the experiment described in Figure 6e. Bubble size represents the proportion of cells in each cluster expressing the marker, and colour scale indicates mean expression level.

**(b–f)** Violin plots showing normalised expression and percentage of cells expressing the indicated markers in (b) all clusters (granularity level 1), (c) cancer cell clusters, (d) B cell clusters, (e) T and NK cell clusters, and (f) myeloid cell clusters. Colours indicate the percentage of cells expressing the marker within each cluster.

**(g–l)** Stacked bar charts showing proportions of cell clusters as percentages of total live cells, for (g) all clusters (granularity level 2), (h) cancer cell clusters, (i) T and NK cell clusters, (j) myeloid cell clusters, (k) B cell clusters, and (l) resident macrophage and MDSC clusters.

**(d)** Dot plot showing the Δarcsinh of pairwise comparisons in the frequency of immune cell clusters identified by spectral flow cytometry in tumours at the endpoint of the experiment described in (e). n = 8 mice per condition. Only fold changes with FDR ≤ 0.05 are shown.

**(e)** Heatmap showing mean normalised protein expression across immune cell clusters identified by spectral flow cytometry for each experimental condition, relative to the mean expression in tumours at the endpoint of the experiment described in (e). n = 8 mice per condition. Only proteins differentially expressed across at least two conditions (FDR ≤ 0.05) are shown.
